# cPLA_2_α Targeting to Exosomes Connects Nuclear Deformation to LTB_4_-Signaling During Neutrophil Chemotaxis

**DOI:** 10.1101/2025.07.02.662855

**Authors:** Subhash B. Arya, Fatima Jordan-Javed, Kristen Loesel, Yehyun Choi, Samuel P. Collie, Lauren E. Hein, Brendon M. Baker, Euisik Yoon, Carole A. Parent

**Affiliations:** Life Sciences Institute, University of Michigan, Ann Arbor, USA; Department of Cell and Developmental Biology, University of Michigan Medical School, Ann Arbor, USA; Cancer Biology Graduate Program, University of Michigan Medical School, Ann Arbor, USA; Department of Electrical Engineering and Computer Sciences, University of Michigan, Ann Arbor, USA; Cellular and Molecular Biology Graduate Program, University of Michigan Medical School, Ann Arbor, USA; Department of Biomedical Engineering, University of Michigan Medical School, Ann Arbor, USA; Department of Mechanical Engineering, University of Michigan, Ann Arbor, USA; Centre for Nanomedicine, Institute for Basic Sciences, Yonsei University, Seoul, Republic of Korea; Department of Pharmacology, University of Michigan Medical School, Ann Arbor, USA; Rogel Cancer Center, University of Michigan Medical School, Ann Arbor, USA

**Keywords:** cPLA_2_α, nuclear envelope, Leukotriene B_4_, exosomes, mechanotransduction, chemotaxis

## Abstract

Efficient neutrophil chemotaxis requires the integration of mechanical forces and lipid-mediated signaling. While the signaling lipid leukotriene B4 (LTB_4_) reinforces cellular polarity, how mechanical cues regulate its production remains unclear. We now show that cytosolic phospholipase A2α (cPLA₂α), which is essential for the synthesis of LTB_4_, functions as a nuclear curvosensor. cPLA₂α responds to nuclear constrictions by localizing to ceramide-rich inner nuclear membrane microdomains and incorporating onto the exofacial surface of nuclear envelope-derived exosomes. This unique topology enables localized LTB_4_ synthesis, which promotes myosin light chain II phosphorylation, and sustains polarity and directional persistence after constriction. In neutrophils squeezing through small constrictions, loss of cPLA₂α impairs nuclear curvature sensing, exosomal LTB_4_ production, and post-constriction motility. These findings uncover a cPLA_2_-dependent mechano-chemical axis linking nuclear architecture to chemotactic efficiency and offering new strategies to modulate inflammatory responses.

## INTRODUCTION

The four cardinal signs of inflammation are tumor (swelling), rubor (redness), dolor (pain), and calor (heat) (*1*). While redness, heat, and pain are mediated by chemicals such as histamine and prostaglandins, swelling results from the accumulation of fluids and immune cells in the affected tissue (*2*). Neutrophils are the first innate immune cells recruited to inflamed or infected sites in response to damage-associated molecular patterns (DAMPs), such as the chemoattractant N-formylmethionine-leucyl-phenylalanine (fMLF), in a process known as chemotaxis (*3*). *In vivo*, neutrophils engage in an actomyosin-driven, two-step “search and run” response to primary chemoattractants(*4*, *5*). This directional migration is amplified by neutrophil-derived leukotriene B4 (LTB_4_), a secondary chemoattractant that is essential for integrin-independent, rapid recruitment of neutrophils throughout injured tissues(*6*).

As the largest and stiffest organelle in the cell, the nucleus is a key regulator of inside-out signaling required for successful mechanotransduction – a process that converts mechanical input into biochemical signaling and gene expression changes (*7*). Changes in nuclear shape, size, and envelope composition influence how cells respond to mechanical forces, a process known as mechanosensing, through nuclear mechanotransduction. For example, depletion of the nucleoskeleton protein, Lamin A/C (LMNA/C), promotes translocation of cytosolic phospholipase A2 alpha (cPLA₂α) to the nuclear envelope (NE) upon swelling in HeLa cells and in response to hypotonic environments in zebrafish neutrophils (*8*). Moreover, in confined dendritic cells and zebrafish progenitor cells, cPLA₂α activity promotes the re-distribution of myosin II to the cell cortex, supporting migratory phenotypes (*9*, *10*). Although these results indicate that cPLA_2_α translocation depends on increased nuclear tension, the mechanosensitive role of the nucleus during chemotaxis in human neutrophils – particularly under constricted conditions that mimic tissue infiltration – remains poorly understood.

cPLA2α plays diverse roles in cellular physiology, ranging from initiating lipid signaling cascades to regulating membrane dynamics (*11*). Its primary enzymatic function is the selective hydrolysis of phospholipids at the sn-2 position, releasing arachidonic acid (AA) – the precursor of eicosanoids such as prostaglandins and leukotrienes – and lysophosphatidylcholine (LPC), a cone-shaped lipid that induces positive membrane curvature (*12*). While AA release is central to the biogenesis of inflammatory mediators, LPC production has been implicated in membrane remodeling and curvature generation. These dual outputs allow cPLA₂α to operate at the intersection of signal transduction and membrane mechanics. In addition to its enzymatic activity, cPLA₂α is itself a sensor and promoter of positive membrane curvature (*13*, *14*). Its N-terminal C2 domain facilitates calcium-dependent membrane binding and is sufficient to induce vesiculation both *in vitro* and in cells (*13*). Importantly, this domain exhibits preferential localization to small, highly curved vesicles (∼50 nm), while excluding flatter or larger structures (∼600 nm) (*14*), underscoring its sensitivity to nanoscale curvatures. Such curvature-sensing features are particularly relevant in the nuclei of neutrophils, where extensive NE remodeling is required for the biogenesis of LTB_4_-producing exosomes (*15*). In this context, cPLA₂α may serve a dual role – sensing extreme membrane curvature at budding NE domains and contributing enzymatically to the lipid composition and signaling potential of nascent exosomes.

AA release from membrane phospholipids is central to eicosanoid metabolism, serving as the precursor for both prostaglandins and leukotrienes. Although neutrophils express cyclooxygenases – the enzymes required for prostaglandin synthesis on lipid bodies – they typically contain few or no lipid droplets per cell (*16*) and produce minimal amounts of prostaglandins (*17*). As a result, leukotriene biosynthesis, particularly LTB_4_, predominates as the principal eicosanoid output in neutrophils. Upon chemoattractant stimulation, G protein-coupled receptor (GPCR) signaling induces a rise in intracellular calcium, which activates cPLA_2_α and releases AA (*18*). The endoplasmic reticulum (ER)/NE-resident, 5-lipoxygenase activating protein (FLAP) then shuttles AA to 5-lipoxygenase (5LO), which sequentially converts it to 5-hydroxyeicosatetraenoic acid (5-HETE) and leukotriene A_4_ (LTA_4_); LTA_4_ hydrolase (LTA_4_H) subsequently generates LTB_4_ (*19*). During chemotaxis, this entire LTB_4_-synthesizing ensemble – 5LO, FLAP, and LTA_4_H – is loaded onto NE buds originating from neutral-sphingomyelinase (nSMase)-generated lipid microdomains on the NE (*20*). These buds mature into NE-derived multivesicular bodies (NE-MVBs) that fuse with the plasma membrane, releasing FLAP-positive, CD63-low exosomes enriched in LTB_4_ (*15*). Extracellular LTB_4_ acts in an autocrine and paracrine fashion by binding to specific GPCRs, to amplify the recruitment of additional neutrophils, sharpen the front-to-rear polarity of cells, and sustain high migratory persistence (*21*). Central to this amplification loop is the phosphorylation of myosin light chain II (MLC II), which reinforces rearward actomyosin contractility (*22*). In addition, LTB_4_ signaling simultaneously engages additional pathways that coordinate actin polymerization, integrin activation, and directional sensing, thereby ensuring robust collective chemotaxis (*23*).

Neutrophils respond to both chemical gradients and physical cues such as matrix stiffness, topography, and architecture (*24*). While components like integrins, TRPV/PIEZO channels, and the cytoskeleton have been implicated in mechanical responses, how mechanical and chemical signals are integrated in fast-migrating cells like neutrophils remains largely undefined (*25*). Notably, inhibition of cPLA₂α in human neutrophils reduces LTB_4_ production and impairs transepithelial migration (*26*). Yet the mechanistic underpinnings linking cPLA₂α activity to exosome biogenesis and chemotaxis have remained elusive. In this study, we employed a novel Constricted Chemotaxis Chamber (C3) that introduces a single, physiologically relevant nuclear deformation during migration to interrogate how mechanical stress on the nucleus regulates signaling outputs. We identify a mechanism by which nuclear squeezing promotes cPLA₂α packaging onto the NE-derived exosomes, and subsequent LTB_4_-dependent reinforcement of neutrophil chemotaxis.

## RESULTS

### Neutrophil nuclear morphology is regulated by cPLA_2_α

To directly assess the role of cPLA2α in nuclear mechanotransduction during neutrophil chemotaxis, we used human promyelocytic leukemia-derived HL60 cells, which can be readily differentiated into neutrophil-like cells in the presence of DMSO (*27*), and generated *cPLA_2_α* knockout (KO) cells using CRISPR-cas9 (**Fig. S1A**). To assess cPLA_2_α-specific effects, we exogenously expressed GFP-cPLA_2_α in *cPLA_2_α* KO (**Fig. S1B**). Surface expression of CD11b levels on differentiated HL60 (dHL60) neutrophils confirmed comparable levels of differentiation across all cell lines – Scr, *cPLA_2_α* KO, and GFP-cPLA_2_α (**Fig. S1C**). We first measured the expression and localization of the nuclear structural proteins, LMNA/C and lamin B receptor (LBR), both implicated in neutrophil nuclear lobulation (*28*). While LBR expression remained unchanged across cell lines, *cPLA_2_α* KO (dHL60) neutrophils exhibited a significant upregulation of LMNA/C compared to Scr dHL60 neutrophils (**Fig. 1A**). Similar results were obtained by quantifying LMNA/C immunofluorescence signals in chemotaxing cells under agarose (**Fig. 1B**). Although dHL60 neutrophils lack distinct nuclear lobes observed in human PolyMorphonuclear Neutrophils (PMNs), they display a complex, folded nuclear morphology characterized by multiple NE invaginations/folds enriched with the inner nuclear membrane (INM)-resident protein LBR (*28*, *29*) (**Fig. 1C**, Scr cells). In contrast, *cPLA_2_α* KO cells exhibited a rounder and less folded nuclear morphology (**Fig.1C**). Quantification of LBR-positive NE folds/invaginations and nuclear surface complexity (form factor) – measured using CellProfiler-based object segmentation – revealed a reduction in LBR-positive invaginations and an increase in nuclear form factor in the KO cells relative to the Scr controls (**Fig. 1D-E**). Notably, re-expression of GFP-cPLA_2_α in KO cells restored the nuclear lobulation and surface complexity to levels comparable to Scr cells (**Fig. 1C-E, Movie S1**). Together, these findings demonstrate that cPLA_2_α regulates LMNA/C expression and is essential for maintaining the characteristic nuclear morphology of neutrophils.

**Fig. 1.**
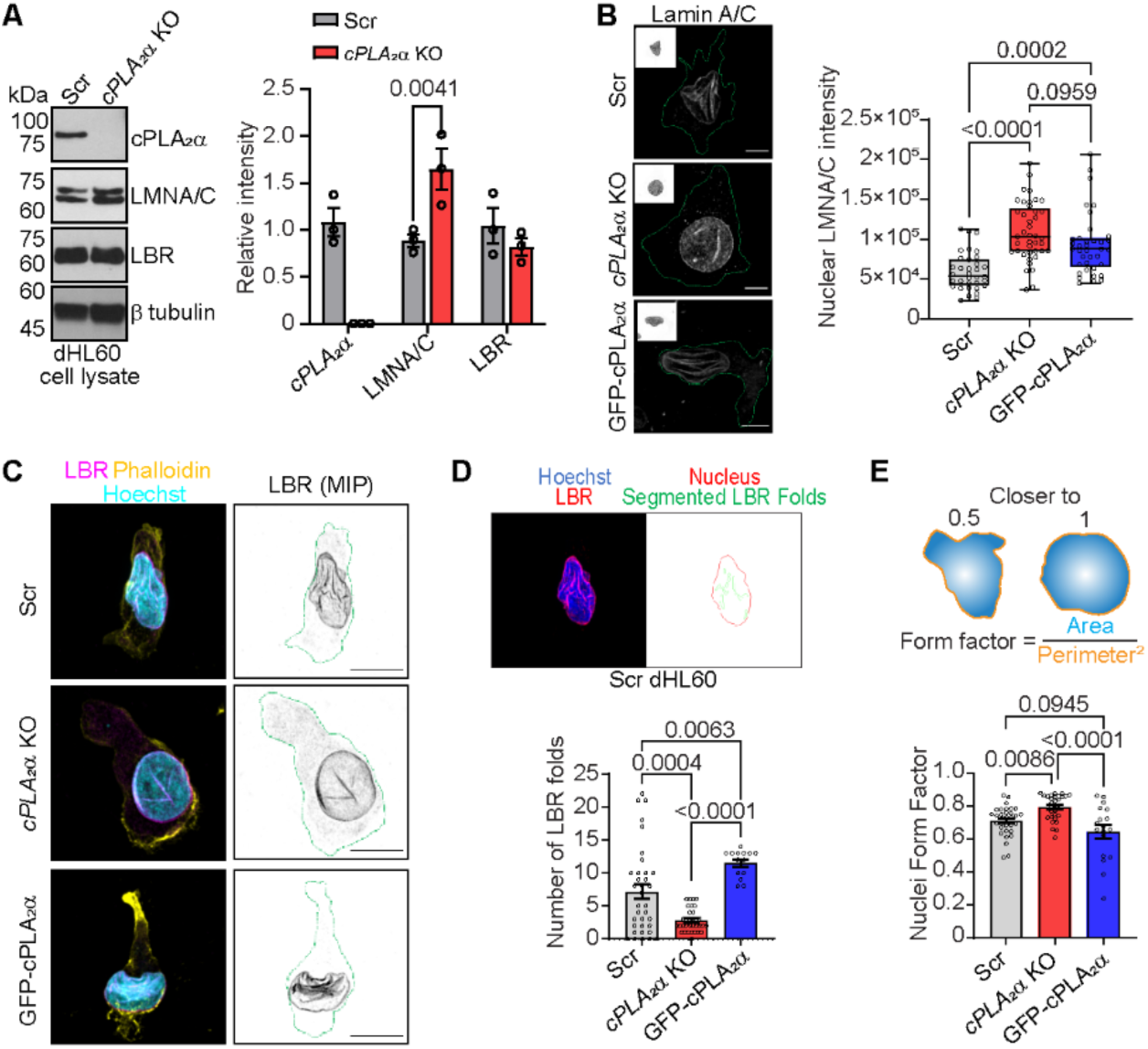
Effect of cPLA_2_α on nuclear morphology in dHL60 neutrophils. **(A-B)** (A) Immunoblots and (B) airyscan microscopy images of dHL60 neutrophils along with respective quantification graph showing the level of LMNA in the cell lysates (A) and nucleus (B). At least 30 data points (B) pooled from 3 independent experiments (A&B) are plotted as mean ± s.e.m. The scale is 5 µm in panel B. **(C)** Airyscan microscopy images of dHL60 neutrophils migrating towards fMLF under agarose, fixed, and immunostained for LBR (magenta), phalloidin (yellow), and Hoechst (blue). **(D-E)** Graphs showing the effect of cPLA_2_α on the number of NE folds and nuclei form factor, plotted as datapoints (circles) pooled from 3 independent experiments. The top panels show nuclear outlines and LBR folds (D) as segmented using CellProfiler, and nuclear morphology described by form factor (E). *P* values determined using ordinary one-way ANOVA (B, D, & E) and two-way ANOVA (A) are shown.

### Neutrophils lacking cPLA_2_α exhibit defective nuclear mechanosensitivity

Since both the LMNA/C (*30*) and cPLA_2_α (*8*, *9*) are involved in regulating cellular and nuclear mechanosensing in cancer cells, necrotic epithelial cells, and dendritic cells, we next evaluated the role of cPLA_2_α on cellular mechanosensing and nuclear mechanotransduction in migrating neutrophils. Efficient nuclear mechanotransduction is reflected by proportional changes in nuclear shape relative to cell shape in response to mechanical cues (*31*). To measure these parameters at the single cell level, we plated fMLF-activated dHL60 neutrophils onto glass coverslips (fibslips) coated with fibrinogen-functionalized synthetic aligned dense fibers (sADF), composed of cell-inert polymer dextran Vinyl Sulfone (DexVS), fabricated by electrospinning (*32s*). We then analyzed cellular and nuclear morphology along the direction of fiber alignment. Within 15 minutes of plating, Scr and GFP-cPLA_2_α cells aligned with the fibslips, and by 30 minutes, their nuclei were embedded and aligned within the ∼15 µm thick sADF mats (**Fig. 2A-B**). Interestingly, while approximately half of the *cPLA_2_α* KO neutrophils aligned with the sADF within 30 minutes (**Fig. 2A-C, G, I**), significantly fewer oriented their nuclei with the fiber direction or intercalated within fibers compared to Scr or GFP-cPLA_2_α cells (**Fig. 2B&D**). Although nuclear volume was comparable across all cell lines (**Fig. 2E**), *cPLA_2_α* KO cells exhibited increased nuclear height with a concomitant decline in nuclear elongation relative to Scr and GFP-cPLA_2_α neutrophils within sADF-aligned populations (**Fig. 2F-H, Movie S2**). Consistent with this, we observed a marked decrease in the correlation between cellular and nuclear elongation in fMLF-activated *cPLA_2_α* KO neutrophils compared to Scr and GFP-cPLA_2_α neutrophils on sADF (**Fig. 2J**). These findings suggest that in *cPLA_2_α* KO neutrophils, increased nuclear stiffness – driven by elevated LMNA/C expression – impairs the dynamic remodeling of nuclear membrane curvature required for nuclear entry and elongation within the channel-like architecture of dense, aligned fiber mats.

**Fig. 2.**
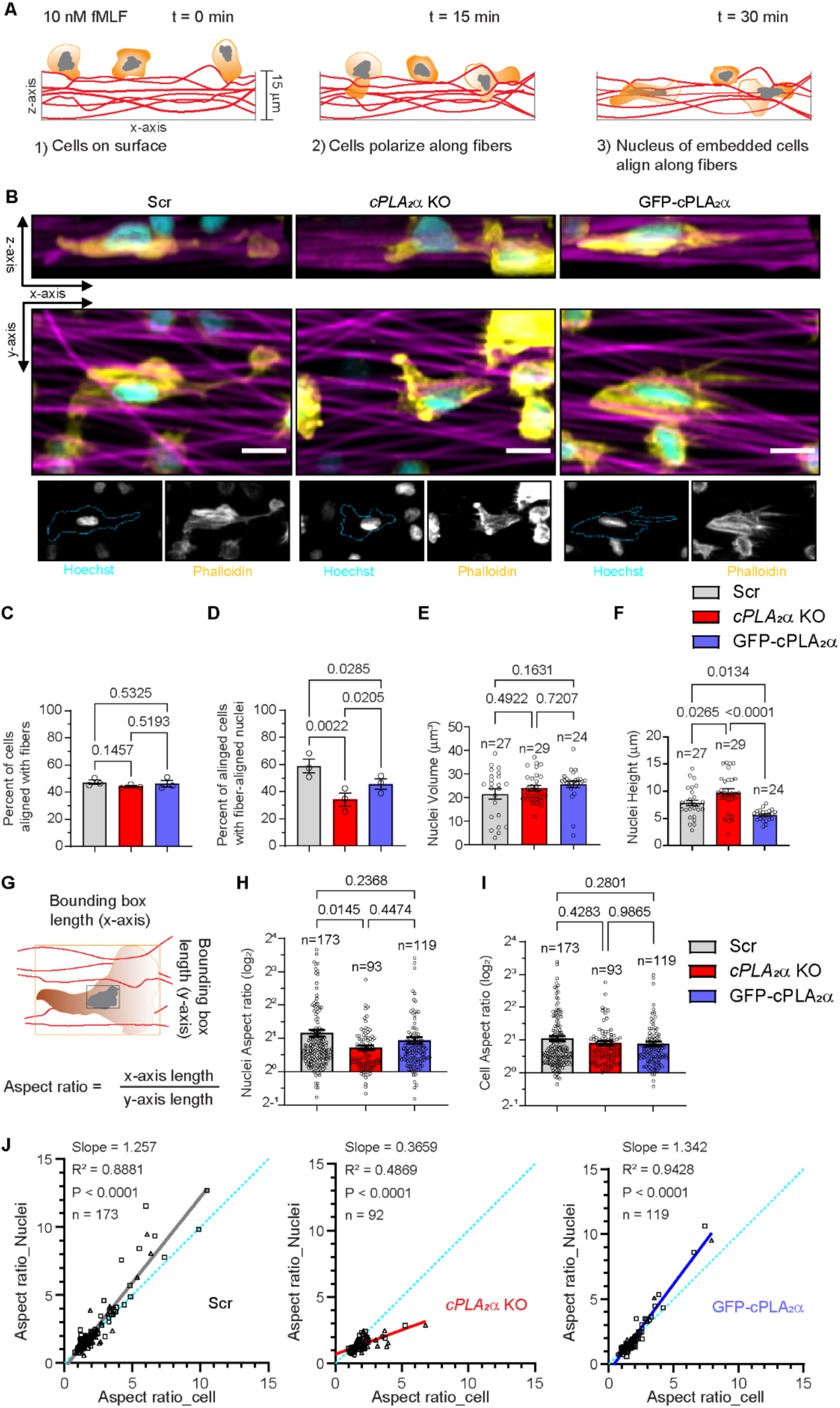
Effect of cPLA_2_α on cellular mechanosensing and nuclear mechanotransduction in activated dHL60 neutrophils. **(A)** Schematic depicting the behavior of neutrophils and their nuclei when plated on sADF fibslips in the presence of fMLF. **(B)** Confocal microscopy images of dHL60 neutrophils showing the shape of cells (phalloidin, yellow) and their nuclei (Hoechst, cyan) on aligned fibers (magenta), along the xy– and xz-axis, 30 min post fMLF treatment. **(C-D)** Graphs plotted as mean ± s.e.m. showing the percent of cells aligned with fibers (C) and the percent of aligned cells with fiber-aligned nuclei (D). N=3. **(E-F)** Graphs plotted as mean ± s.e.m. of data points pooled from 3 independent experiments, showing the nuclear volume (E) and the height of nuclei (F) within all aligned cells. **(G)** The approach used to calculate the cell and nuclei aspect ratio. The orange outline denotes the cell bounding box, and the black outline denotes the nuclear bounding box. Wavy red lines denote aligned microfibers. **(H-I)** Graphs plotted as mean ± s.e.m. of data points pooled from 3 independent experiments, showing the changes in the nuclear (H) and cellular (I) aspect ratio in *cPLA_2_α* KO and GFP-cPLA_2_α cells. *P* values determined using RM one-way ANOVA (C-D) and ordinary one-way ANOVA (E-F, H-I) are shown. **(J)** Graphs showing the correlation of cell and nuclei aspect ratio in dHL60 neutrophils plated on aligned fibers and activated with fMLF for 30 min.

### Nuclear constriction promotes cPLA_2_α-dependent increases in directionality during neutrophil chemotaxis

To investigate how neutrophils specifically respond to nuclear constrictions during chemotaxis, we developed the Constricted Chemotaxis Chamber (C^3^), a microfluidic device designed to expose chemotaxing cells to a single, transient constriction. The C^3^ device is composed of polydimethylsiloxane (PDMS) bonded to a glass coverslip via plasma activation and features diametrically opposed inlets and outlets for cell and chemoattractant loading, each 100 µm high (**Fig. 3A**). The migration channels are 5 µm in height, 300 µm in length, and contain mechanical confinement pillars of various gap sizes positioned 100 µm from the cell inlet (**Fig. 3A**, zoomed inset). We selected a cross-shaped constriction pillar (20 µm across) over conventional cylindrical posts (*33*, *34*) as the cross geometry produced higher curvature constrictions suited for our assay, whereas cylindrical columns frequently resulted in undesired cell wrapping during migration (data not shown). This design enables transient entrapment of migrating neutrophils in a cup-like space between two constriction pillars, compelling them to squeeze through defined constriction gaps (**Fig. 3A**, zoomed inset). Given the average nuclear diameter of dHL60 neutrophils (∼4 µm; **Fig. 2E**), we selected constriction widths of 5– and 3-micron to represent conditions of unrestricted and restricted nuclear passage and nuclear squeezing, respectively. Finally, using Alexa Fluor 488-tagged fMLF, we verified that the chemoattractant gradient establishes within 15 minutes and remains stable over 60 minutes without external flow control (**Fig. 3B)**. Thus, the C^3^ device allows direct visualization and quantification of nuclear squeezing and its impact on neutrophil chemotaxis (**Fig. 3C**).

**Fig. 3.**
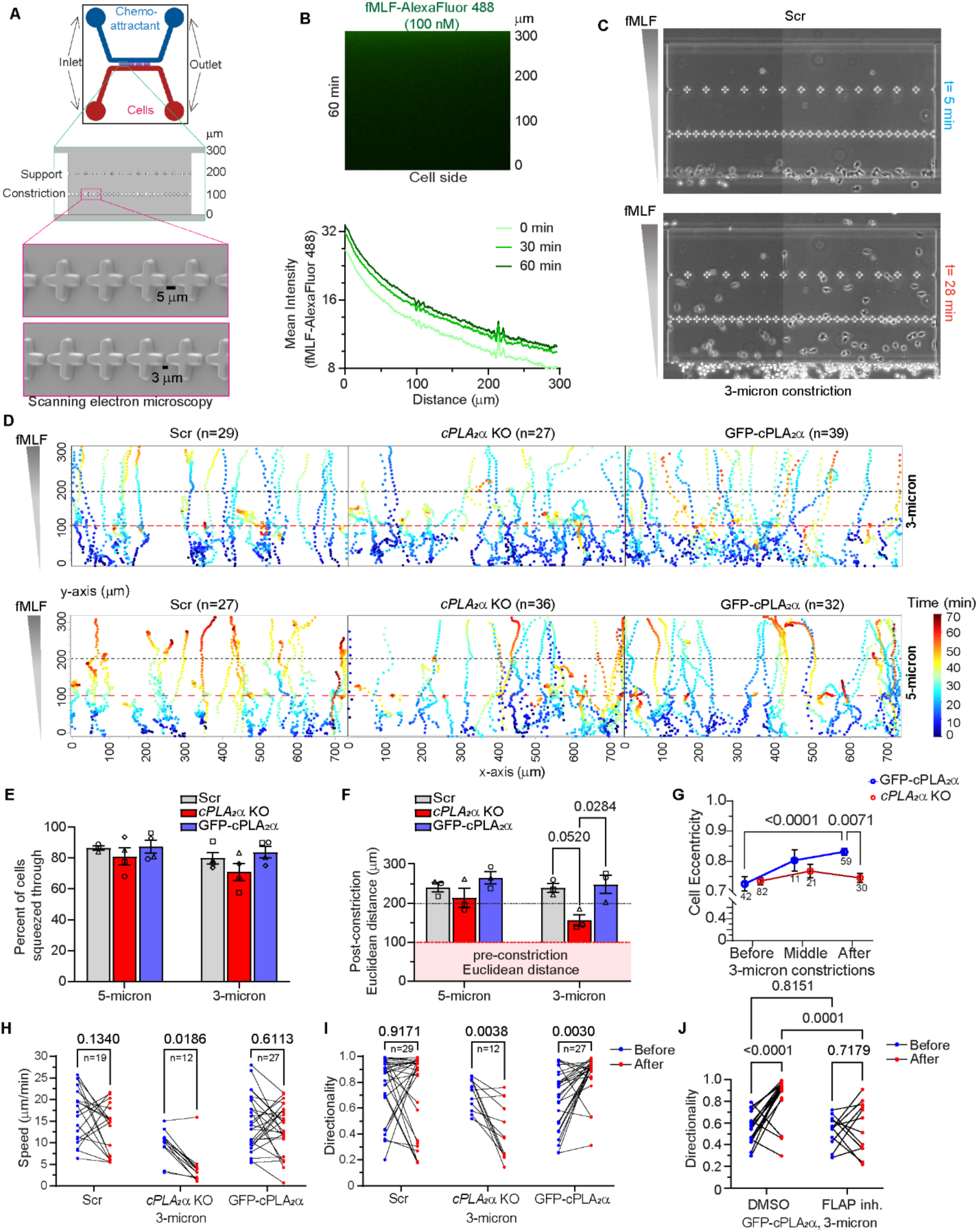
Effect of cPLA_2_α on dHL60 neutrophils chemotaxing through transient constrictions. **(A)** Image of the C^3^ showing the cell and chemoattractant inlet/outlet and the migration chamber. The zoomed inset (green) shows the migration chamber, and the zoomed inset (magenta) shows SEM images of the 3– and 5-micron constrictions. **(B)** Image (top) and graph (bottom) showing the diffusion rate and gradient stability of fMLF-Alexa Flour 488 in C^3^. **(C)** Phase contrast images of Scr dHL60 neutrophils migrating towards fMLF through 3-micron constrictions in C^3^ at different time points. **(D)** Color-coded tracks of individual cells migrating towards the fMLF, through either 5-or 3-micron constrictions. Refer to the temporal color map on the right. The red dashed line indicates the point of constriction 100 µm from the migration start site. The black dashed line indicates the support site. **(E-F)** Graph showing the percentage of cells entering the C^3^ that squeezed through 5-or 3-micron constrictions (E) and the post-constriction Euclidean distance (F). N=3. **(G)** Graph plotted as mean ± s.e.m. showing the change in cell eccentricity in response to constrictions during chemotaxis. N=3. **(H-I)** Before-after graph showing the change in speed (H) and directionality (I) of chemotaxing cells post 3-micron constrictions. (**J)** Before-after graph showing the change in the directionality of chemotaxing GFP-cPLA_2_α expressing cells treated with either DMSO or FLAP inhibitor in 3-micron constrictions. *P* values determined using two-way ANOVA are shown.

Analysis of Hoechst-stained dHL60 neutrophils revealed that at least 80% of Scr, GFP-cPLA_2_α, and, surprisingly, *cPLA_2_α* KO neutrophils successfully migrated through 3– and 5-micron constrictions within one hour (**Fig. 3D, E**). Using the Image J ‘TrackMate’ plugin for cell tracking and MATLAB for speed and directionality analysis (see Methods), we found that most Scr and GFP-cPLA_2_α dHL60 neutrophils migrated ∼150 µm beyond the constriction, regardless of constriction width (**Fig. 3D&F**). We reason that the cell arrest near 250 µm from the inlet likely reflects saturating chemoattractant levels near the fMLF outlet (300 µm from the cell inlet). Interestingly, while *cPLA_2_α* KO cells performed similarly to Scr and GFP-cPLA_2_α cells in the 5-micron C^3^ device, they migrated significantly shorter distances after traversing 3-micron constrictions (**Fig. 3D&F; Movie S3**). This post-constriction impairment correlated with loss of cell polarity in *cPLA_2_α* KO cells, in contrast to the enhanced polarity observed in GFP-cPLA_2_α cells (**Fig. 3G**). Moreover, unlike Scr and GFP-cPLA_2_α neutrophils, *cPLA_2_α* KO cells failed to increase their speed after 3-micron constrictions and exhibited reduced directionality (**Fig. 3H&I, S2A&B**). Notably, GFP-cPLA_2_α cells, which express elevated cPLA_2_α levels compared to Scr controls (**Fig. S1B**), showed a significant post-constriction increase in directionality (**Fig. 3I, S2B**). To determine whether this response depends on leukotriene signaling, we pretreated GFP-cPLA_2_α cells with MK866, a FLAP inhibitor that blocks LTB_4_ production (*35*). Strikingly, MK886 treatment abolished the post-constriction increase in directionality observed in untreated GFP-cPLA_2_α cells (**Fig. 3J**). Together, these results suggest that cPLA_2_α levels modulate neutrophil chemotactic performance in response to nuclear squeezing, with higher expression promoting directional persistence in an LTB_4_-dependent manner.

While Scr dHL60 neutrophils initially migrated in an amoeboid pattern through the initial one-third of the C^3^ chamber (before the constrictions), they transitioned into a fan-shaped, keratocyte-like morphology as they approached higher fMLF concentrations – particularly beyond the 3-micron constrictions (**Fig. 4A, top panel**). Keratocyte-like cells, defined by a morphology showing a major axis perpendicular to the direction of migration, and exhibit a smoother cell membrane compared to amoeboid cells (*36*, *37*). Based on this information, we used CellProfiler-based object segmentation (**Fig. 4B**) to classify migrating neutrophils as keratocyte-like if their orientation exceeded 45°, eccentricity >0.75, and solidity >0.9. Notably, we found that cPLA_2_α depletion reduced this amoeboid-to-keratocyte-like transition by nearly 50% (**Fig. 4A&C**). Accompanying this defect, we observed persistent uropod extension in *cPLA_2_α* KO cells that remained polarized post-constriction, while most others lost polarity (**Movie S3**). Together, these results indicate that cPLA_2_α is critical for the morphological and migratory transitions required for directional persistence in neutrophils undergoing nuclear squeezing.

**Fig. 4:**
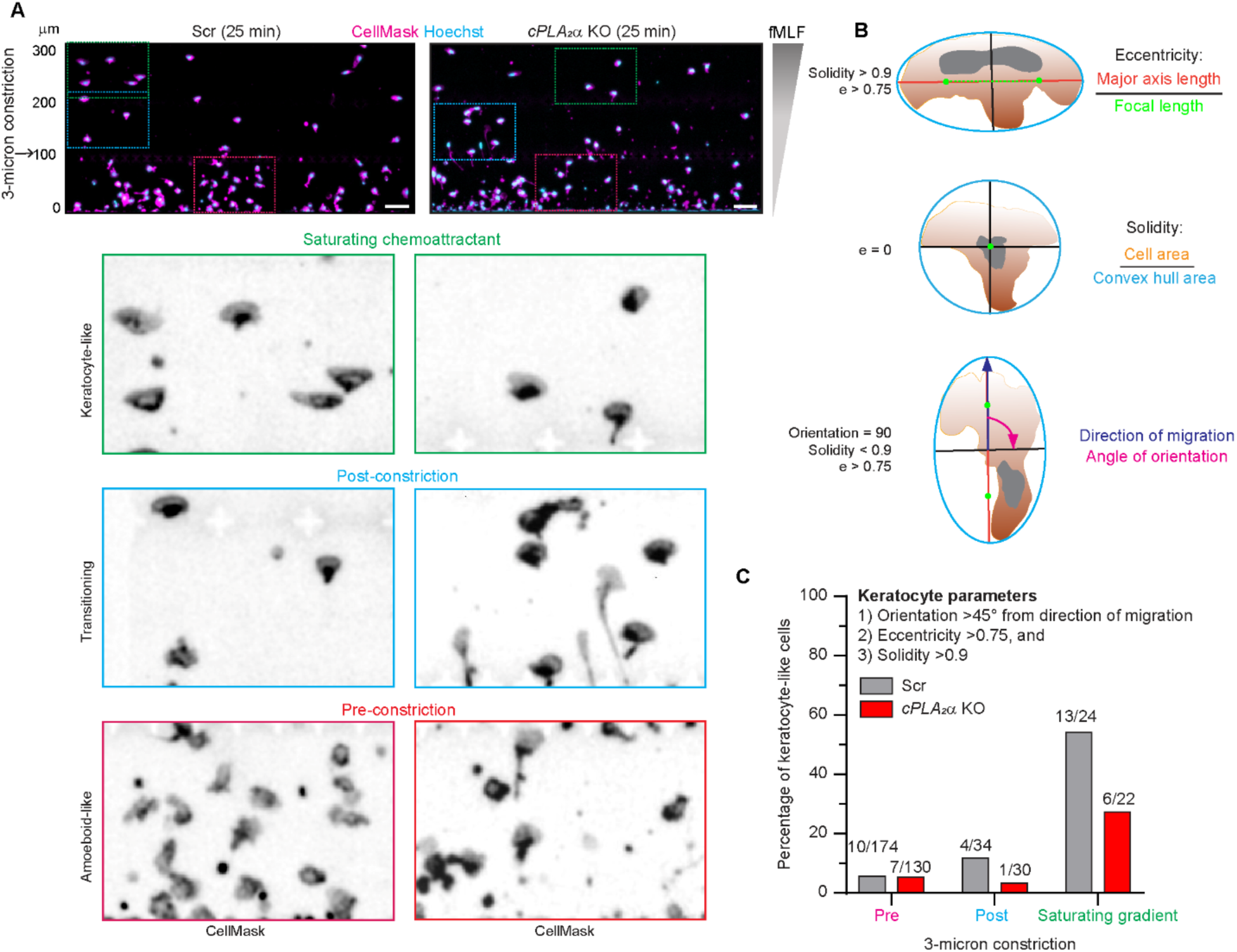
Loss of cPLA2α inhibits amoeboid-to-keratocyte-like migratory transition in response to nuclear squeezing. **(A)** Microscopy images of dHL60 cells stained with CellMask (PM, magenta) and Hoechst 33342 (nucleus, cyan) 25 min after the start of chemotaxis in 3-micron C^3^ devices, showing the morphological transitions from amoeboid-like to keratocyte-like migration mode (red-cyan-green insets). The scale is 10 μm, and graph **(B)** Schematic showing the metrics used for measuring various cellular parameters. **(C)** Graph showing the percentage of keratocytes in Scr vs. *cPLA_2_α* KO cells, and the classifier used to ascertain keratocyte-like morphology.

### Nuclear constriction drives cPLA_2_α-dependent MLC II phosphorylation during neutrophil chemotaxis

Persistent migration requires robust actomyosin contractility, particularly at the cell rear (*4*). We previously demonstrated that inhibition of LTB_4_ biosynthesis in fMLF-stimulated neutrophils markedly reduces phosphorylation of myosin light chain II (MLC II) and impairs chemotaxis (*21*, *22*). Given that migration through 3-micron constrictions induces a morphological transition from an amoeboid-to a keratocyte-like phenotype (**Fig. 4**) and enhances directional persistence (**Fig. 3I**), we next investigated whether this transition involves changes in the cellular levels of myosin and its cortical redistribution.

Using object-based segmentation, we quantified levels and localization of phosphorylated MLC II (pMLC II) in *cPLA_2_α* KO and GFP-cPLA_2_α neutrophils before, during, and after passage through 5– and 3-micron constrictions. We observed a significant increase in intracellular pMLC II intensity specifically in GFP-cPLA_2_α cells during and after transit through 3-micron constrictions (**Fig. 5A&B**). Moreover, the cortex-to-cytosol pMLC II ratio increased exclusively in GFP-cPLA_2_α cells navigating through 3-micron constrictions (**Fig. 5C**), coinciding with enhanced cell polarity (**Fig. 3G**). These findings indicate that cPLA_2_α promotes actomyosin contractility in response to nuclear squeezing, enabling transition from an amoeboid to persistent, keratocyte-like migration.

**Fig. 5.**
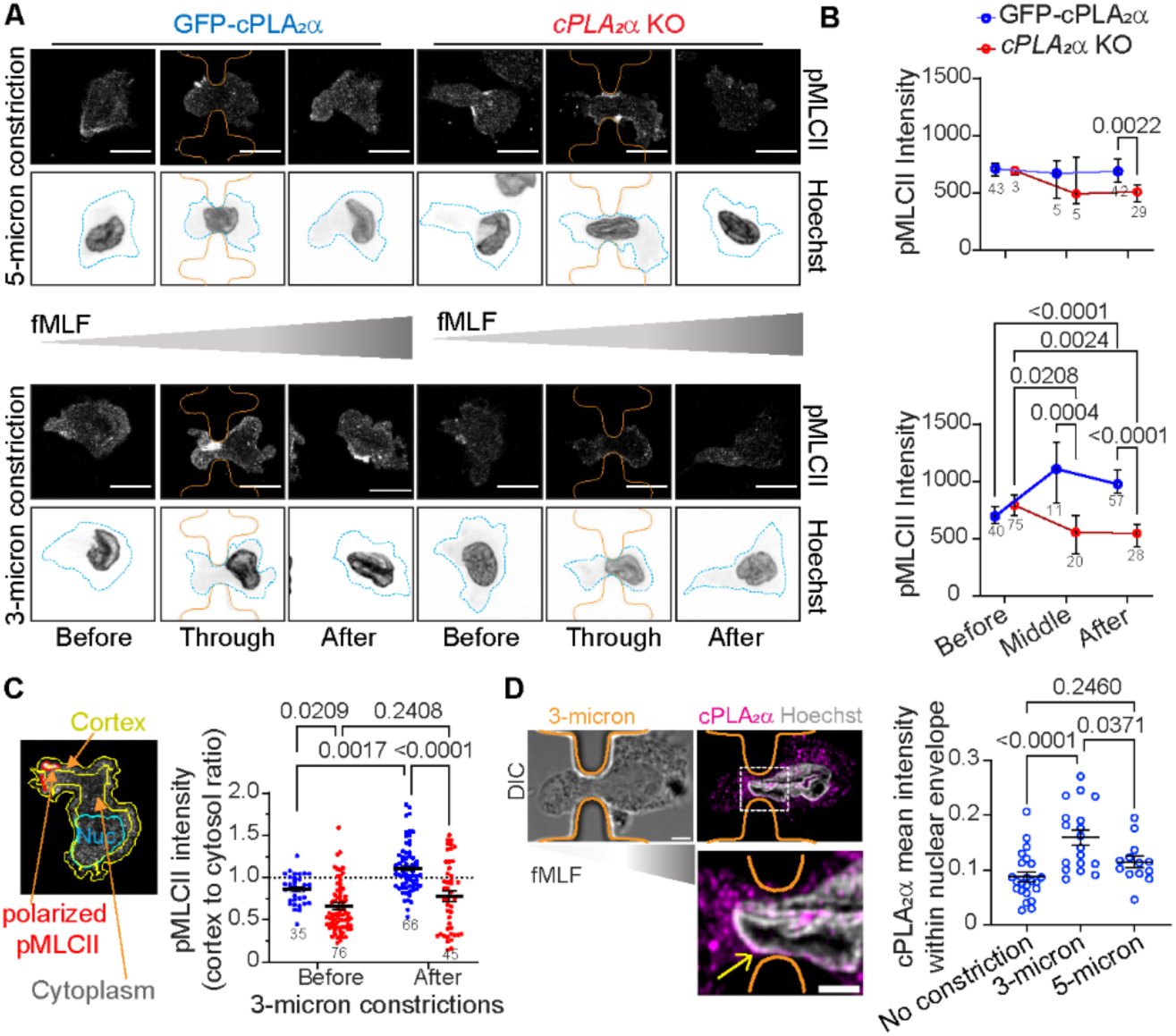
Effect of cPLA_2_α on post-constriction MLC II phosphorylation and cell polarization in dHL60 neutrophils. **(A)** Confocal microscopy images of dHL60 neutrophils chemotaxing towards fMLF, fixed and immunostained for pMLC II and Hoechst 33342 (nuclei). Blue outline indicates the cell periphery, and orange outline indicates constriction pillars. The scale is 10 µm. **(B)** Graphs plotted as mean ± s.e.m., showing the change in total pMLC II intensity within the cell before, during, and after constrictions. N=3. **(C)** Representative microscopy image (left) showing the cortex (yellow), nucleus (cyan), and polarized pMLC II (red) outlines determined using CellProfiler-based object segmentation. The scatter dot plot (right) shows the cortex-to-cytosol pMLC II intensity ratio in the chemotaxing cells before and after 3-micron constrictions. *P* values determined using two-way ANOVA are shown. The number of data points pooled from 3 independent experiments is mentioned on each graph. **(D)** Airyscan microscopy images (left) of a GFP-cPLA_2_α dHL60 neutrophil squeezing through a 3-micron constriction, immunostained for cPLA_2_α (magenta) and Hoechst (gray), along with the phase contrast image. Orange outlines the constriction pillars, and the yellow arrow points to the NE enrichment of cPLA_2_α at the constriction. Scale is 3 μm, it’s 2 μm in the zoomed inset. Scatter dot plot (right) showing the changes in cPLA_2_α levels at the NE before and after 3– and 5-micron constrictions. Datapoints (circles) from 3 independent experiments were pooled, and *P* values determined using two-way ANOVA are presented.

We envision that tight nuclear constriction activates cPLA_2_α, triggering LTB_4_ production and downstream MLC II phosphorylation – key events that enhance post-constriction motility and directional persistence during neutrophil chemotaxis. Zebrafish cPLA_2_α, a predominantly nuclear-localized protein, has been shown to translocate from the nucleus to the nuclear envelope (*8*, *38*). Consistent with previously reported nucleo-cytosolic distribution of cPLA_2_α (*39*, *40*), we found that in chemotaxing GFP-cPLA_2_α neutrophils, ∼30% of the GFP-cPLA_2_α signal was nuclear, with ∼20% localized to the nucleoplasm and ∼10% enriched at the NE (**Fig. S3A&B**). Notably, GFP-cPLA_2_α further accumulated at the NE in neutrophils migrating through 3-micron (but not 5-micron) constrictions, coinciding with increased pMLC II levels (**Fig. 5D**). Using 4x expansion microscopy to enhance NE resolution, we found that in chemotaxing neutrophils, GFP-cPLA_2_α localized to LBR-positive INM, but not to calnexin-positive outer nuclear membranes (ONM) (**Fig. 6A**). These results suggest that nuclear squeezing promotes the recruitment of cPLA_2_α to INM.

**Fig. 6.**
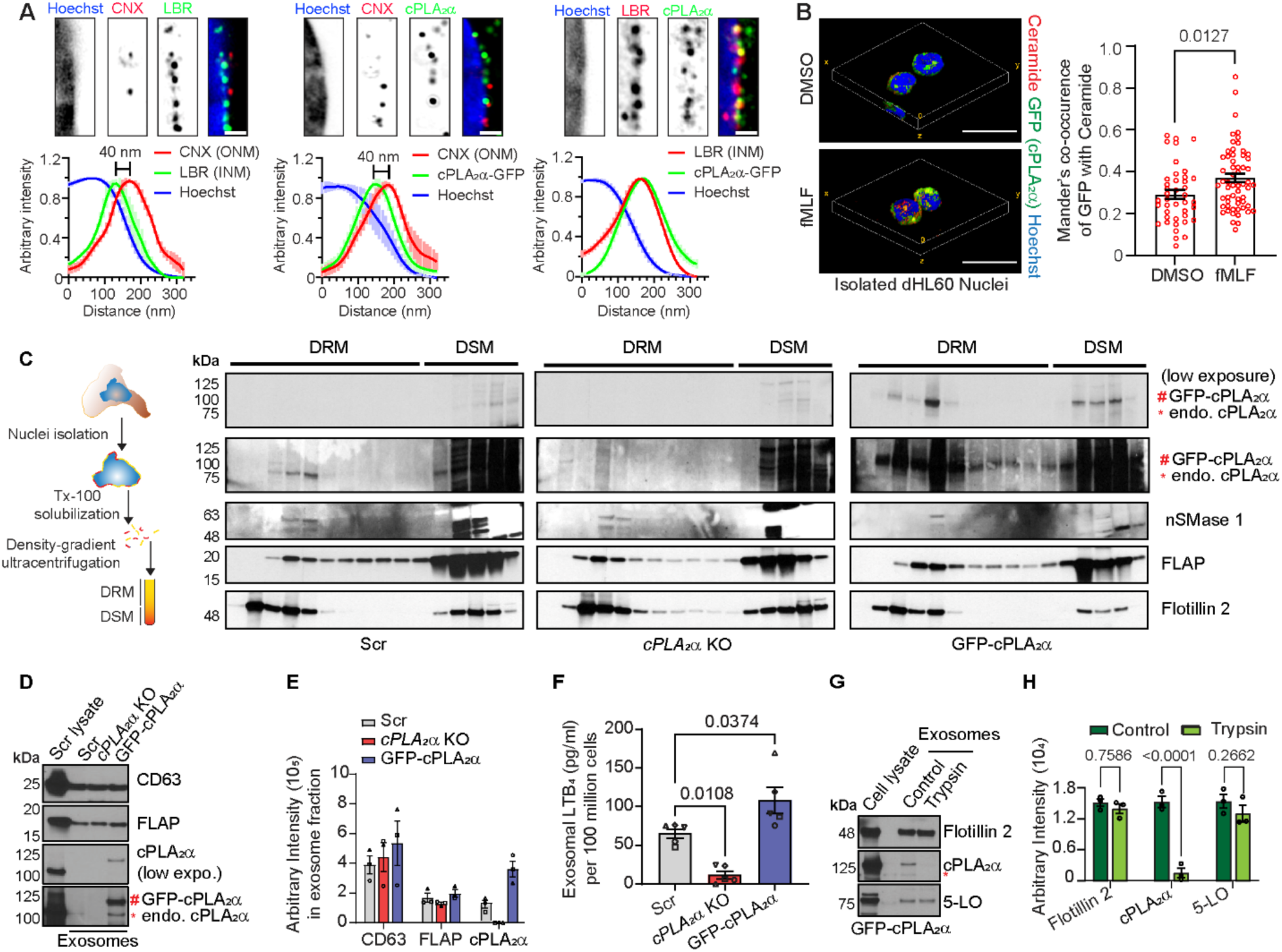
cPLA_2_α is enriched at INM microdomains and on the exofacial surface of exosomes in activated dHL60 neutrophils. **(A)** Four-fold expansion microscopy images of chemotaxing dHL60 neutrophils expressing GFP-cPLA_2_α and immunostained with the indicated antibodies. The scale is 300 nm. Corresponding histograms below each panel show fluorescence intensity profiles averaged across multiple images, plotted as mean ± s.e.m.; solid lines represent the mean, and shaded bars indicate the standard error. N=3. **(B)** 3D volumetric view (left) of fixed and immunostained nuclei isolated from dHL60 neutrophils showing the distribution of GFP-cPLA_2_α (green), ceramide (red), and Hoechst 33342 (blue), quantified and plotted in a graph (right) showing Mander’s co-occurrence. Datapoints (red circles) are plotted as mean ± s.e.m., and *P* values determined using Mann Mann-Whitney test are shown. N=3. The scale is 10 µm. **(C)** Schematic (left) showing the steps involved in the isolation of NE-membrane microdomains, and immunoblots (right) showing the cPLA_2_α, FLAP, and Flotillin 2 distribution in DRM and DSM fractions of NE obtained from activated cells. N=4. **(D-E)** Immunoblot (D) and graph (E) plotted as mean ± s.e.m., showing the levels of CD63, FLAP, and cPLA_2_α in purified exosomes obtained from various cell lines. N=3. **(F)** Graph showing the levels of LTB_4_ within the exosomes purified from various cell lines upon fMLF activation. N=5. **(G-H)** Immunoblots (G) and graph (H) plotted as mean ± s.e.m., showing the levels of cPLA_2_α, 5LO, and Flotillin 2 in the purified exosomes upon trypsin treatment. N=3. *P* values determined using two-way ANOVA are presented.

### cPLA_2_α is recruited to the exofacial surface of NE-derived exosomes

We previously showed that neutrophils undergoing chemotaxis exhibit NE budding at ceramide-rich, lipid-ordered membrane microdomains (*15*), which requires nSMase activity on the NE (*15*). Moreover, cPLA_2_α-dependent AA release has been proposed to activate nSMase *in vitro* (*41*). We found that in dHL60 neutrophils, GFP-cPLA_2_α colocalized with ceramide-rich membrane microdomains on the NE of isolated nuclei, and Mander’s coefficient analysis revealed increased colocalization upon fMLF stimulation, indicating activation-dependent redistribution (**Fig. 6B, Movie S4**). In addition, both endogenous and GFP-tagged cPLA_2_α were found in lipid-ordered, detergent-resistant membrane (DRM) fractions isolated from the NE of fMLF-stimulated cells (**Fig. 6C**). Importantly, we found that cPLA_2_α is not required for ceramide-rich membrane microdomain formation, as the absence of cPLA_2_α did not alter the enrichment of Flotillin 2, FLAP, or nSMase 1 in these domains (**Fig. 5C, S3C**). Thus, while cPLA_2_α localizes to these lipid microdomains, it is dispensable for their assembly in neutrophils.

Because the LTB_4_ synthesis machinery – 5LO, FLAP, and LTA_4_H – are present in NE-derived LTB_4_-containing exosomes (*15*, *20*), we asked whether cPLA_2_α is also present in these vesicles. Density-gradient ultracentrifugation of fMLF-stimulated dHL60 neutrophil supernatants revealed that cPLA_2_α is associated with 5LO/FLAP-positive exosomes (**Fig. 6D-E**). Although the absence of cPLA_2_α did not affect the recruitment of 5LO or FLAP to exosomes, as expected, LTB_4_ levels were completely abolished in exosomes isolated from *cPLA_2_α* KO cells (**Fig. 6F**). Conversely, GFP–cPLA_2_α neutrophils had elevated exosomal LTB_4_ levels compared to Scr controls, consistent with their higher cPLA_2_α expression (**Fig. 6D–F**).

Given the ability of cPLA_2_α to bind positively curved membranes and the high curvature of the exofacial leaflet of exosomes, we reasoned that cPLA_2_α is present on the outer surface of exosomes. To test this, we treated exosomes purified from fMLF-stimulated GFP-cPLA_2_α dHL60 neutrophils with trypsin. As expected, trypsin did not affect the level of intra-exosomal proteins like Flotillin 2 (*42*) and 5LO, but completely abolished the GFP-cPLA_2_α signal (**Fig. 6G-H**), showing that cPLA_2_α is exposed on the exofacial surface of NE-derived exosomes. Collectively, these findings support a model in which the generation of extreme positive curvature at INM generated during nuclear squeezing enhances nuclear cPLA_2_α enrichment at these sites. This pool of cPLA_2_α becomes incorporated onto the outer surface of highly curved intraluminal vesicles, which are subsequently secreted as NE-derived exosomes enriched in cPLA_2_α and the LTB_4_-synthesis machinery.

### cPLA_2_α-dependent LTB_4_ synthesis in NE-derived exosomes

The catalytic activity and membrane recruitment of cPLA_2_α require elevated intracellular calcium, which can be induced by GPCR signaling (*43*, *44*). Neutrophils migrating under agarose via an adhesion-independent ‘chimneying’ mode depend heavily on actomyosin contractility (*45*, *46*). This prompted us to investigate the localization dynamics of cPLA_2_α in human PMNs, both in suspension and during chemotaxis under agarose. In chemotaxing neutrophils, we observed enrichment of cPLA_2_α at NE budding sites, marked by LBR positivity and exclusion of Hoechst 33342 staining (**Fig. 7A**, red arrow). In contrast, cPLA_2_α was not observed in regions of NE distant from budding sites (**Fig. 7A**, black arrow), as confirmed by line profile analysis. Additionally, cPLA_2_α was detected on LBR-positive cytosolic vesicles that likely originated from NE buds (**Fig. 7B**). Indeed, line profile analysis of NE-derived MVBs in neutrophils co-stained with cPLA_2_α and 5LO (intraluminal vesicle marker) revealed that cPLA_2_α is present on the periphery of 5LO-positive intraluminal structures (**Fig. 7B**).

**Fig. 7.**
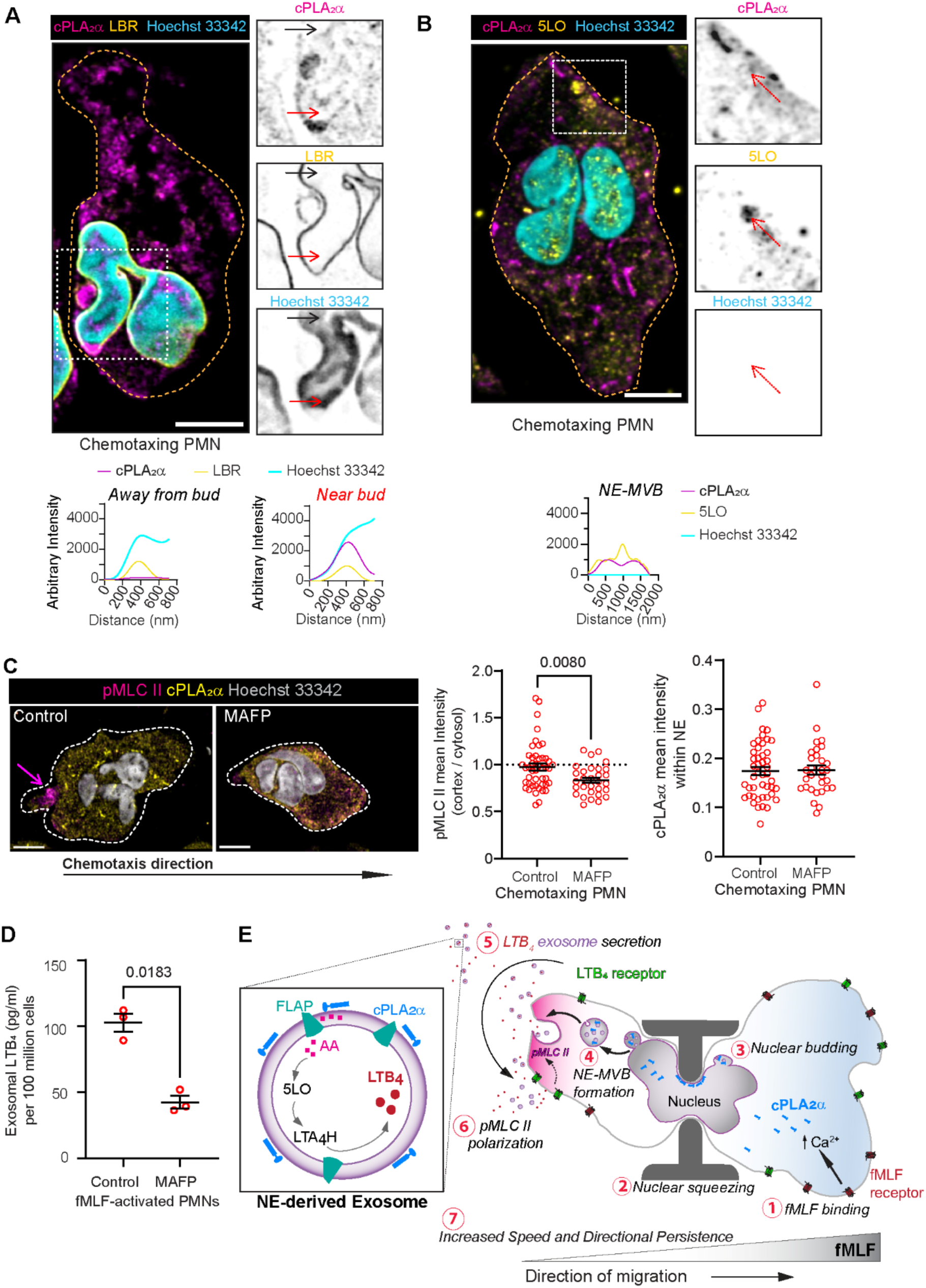
Exosomal cPLA_2_α is enzymatically active and mediates neutrophil chemotaxis. **(A-B)** Airyscan microscopy images of PMNs chemotaxing under agarose, immunostained for cPLA_2_α (magenta) and (A) LBR (yellow) or (B) 5-LO (yellow), and co-stained with Hoechst 33342 (cyan). Orange outlines indicate cell boundaries. The region within the dashed white rectangle is shown on the right as individual inverted grayscale channels. Fluorescence intensity profiles across (A) the red arrow (near NE bud) and black arrow (away from bud) and (B) the red arrow (near cytosolic vesicle) are plotted below the zoomed panels. The scale is 5 μm. **(C)** Confocal microscopy images (left) of PMNs treated with either DMSO or MAFP, chemotaxing under agarose, immunostained for pMLC II (magenta), cPLA_2_α (yellow), and co-stained with Hoechst 33342 (gray); and quantification (right) of cortex to cytosol pMLC II intensity from multiple images plotted as a box whisker plot. P values determined using the Student t-test are presented. Orange outlines indicate cell boundaries. The scale is 5 μm. Red circles on the graph indicate individual datapoints pooled from 3 independent experiments. **(D)** Scatter dot plot showing the levels of LTB_4_ within the exosomes isolated from fMLF activated PMNs and treated either with DMSO or MAFP for 30 min at 37 °C. Datapoints (red circles) representing 3 independent experiments are plotted as mean ± s.e.m. P values determined using the Student t-test are presented. **(E)** Schematic illustrating the proposed model of cPLA₂α recruitment and packaging onto the exofacial surface of NE-derived exosomes in response to nuclear squeezing.

To test whether cPLA_2_α catalytic activity impacts actomyosin contractility, we assessed pMLC II levels in chemotaxing human PMNs treated with either the cPLA_2_α inhibitor MAFP (*47*) or DMSO (control). Although the NE-distribution of cPLA2α remained unchanged, MAFP treatment significantly reduced cortical pMLC II levels, particularly at the cell rear (**Fig. 7C**), indicating that cPLA_2_α-dependent AA production – rather than its subcellular location – is essential for proper actomyosin regulation during chemotaxis. Finally, we found that exosomes isolated from fMLF-stimulated PMNs treated with MAFP have significantly reduced LTB_4_ levels compared to DMSO controls (**Fig. 7D**), showing that exosomal cPLA_2_α is enzymatically active.

Together, findings from this study support a model in which transient nuclear squeezing during migration through tight constrictions induces inward curvature of the INM, driving cPLA_2_α enrichment at budding NE domains. This facilitates the packaging of catalytically active cPLA_2_α into NE-derived exosomes. These exosomes serve as platforms for localized LTB_4_ synthesis, which in turn activate LTB_4_ receptors and promote MLC II phosphorylation. The resulting enhancement of cell polarity and actomyosin contractility sustains persistent chemotactic signaling (**Fig. 7E**).

## DISCUSSION

The LTB_4_ signaling pathway plays a central role in neutrophil extravasation (*23*) and directional migration during inflammation (*6*, *21*). Central to this pathway is cPLA_2_α, which catalyzes the release of AA, the precursor for LTB_4_ biosynthesis (*48*). While the importance of cPLA2α in neutrophil chemotaxis has been shown, the spatial and mechanical contexts in which its activity becomes critical have remained poorly understood. In this study, we reveal that cPLA_2_α not only governs LTB_4_ production but also orchestrates a nuclear mechanotransduction program that enables neutrophils to navigate physical barriers during chemotaxis. Using a custom-engineered chamber, we show that nuclear squeezing through tight constrictions induces NE remodeling and cPLA2α redistribution, promoting LTB_4_ synthesis both intracellularly (*20*) and on secreted exosomes. These exosomes, in turn, reinforce directional migration by activating LTB_4_ signaling and actomyosin contractility. Our findings establish a direct mechanistic link between nuclear deformation, lipid signaling, and chemotactic persistence, highlighting cPLA2α as a key integrator of mechanical and chemical cues during immune cell navigation.

Although the role of cPLA_2_α in sensing nuclear curvature changes under hypotonic stress or mechanical compression is well established (*8*, *49*, *50*), its influence on steady-state nuclear architecture remained unexplored. We found that dHL60 neutrophils lacking cPLA_2_α, despite undergoing normal differentiation, exhibit reduced nuclear curvature and surface complexity. This altered nuclear morphology is accompanied by increased LMNA expression without any change in LBR levels. These observations are consistent with previous reports implicating cPLA_2_α in the regulation of immune-related gene expression in mechanically stimulated mouse dendritic cells (*9*). Furthermore, our findings now show that cPLA_2_α-deficient neutrophils with high LMNA levels exhibit reduced nuclear mechanotransduction, suggesting that cPLA_2_α regulates mechanotransduction, at least in part, by modulating LMNA expression in mature neutrophils. Consistent with the role of LMNA in promoting nuclear stiffness (*51*), we found that *cPLA_2_α* KO neutrophils show impaired global nuclear shape remodeling during migration on sADF matrices. The misalignment of nuclei with the underlying substrate topology observed in *cPLA_2_α* KO neutrophils coincided with a decoupling of cellular and nuclear elongation – indicating impaired nuclear mechanosensitivity. Since we found that nuclear volumes were consistent across all migrating cell types on sADF, we reason that increased LMNA expression and nuclear stiffness, rather than changes in volume and NE unfolding (*10*, *33*), are responsible for the alignment defects observed in polarized *cPLA_2_α* KO neutrophils.

In contrast to aligned collagen fibers of lymph nodes and tumor microenvironments, which are closely mimicked by the organized architecture of sADF (*32*, *52*), the extracellular matrix in inflamed tissue typically consists of disordered collagen networks with variable pore sizes ranging from 0.5 to 10 μm (*53*). This structural heterogeneity makes transient nuclear deformation a frequent occurrence for neutrophils traversing tissues. To model these physiological conditions, we developed a microfluidic device (C^3^) that maintains stable linear chemoattractant gradients while introducing a single, transient constriction during neutrophil chemotaxis. Unlike existing microchannel systems that focus on either mechanotransduction (unidirectional, no choice) or chemotaxis alone, our device enables two-dimensional (bidirectional choice) migration, enabling simultaneous assessment of how transient nuclear squeezing affects directionality, speed, and polarization. Using the C^3^ device, we observed no significant differences in migration distance, speed, or directionality among Scr, *cPLA_2_α* KO, and GFP-cPLA_2_α neutrophils migrating through 5-micron constrictions. Given that the average nuclear diameter of dHL60 neutrophils is ∼4 μm, 5-micron constrictions do not impose substantial nuclear deformation and thus recapitulate behaviors observed in cells chemotaxing under agarose. Notably, all three cell types showed comparable efficiency in crossing 3-micron constrictions, indicating that neither cPLA_2_α deficiency nor elevated LMNA levels impair neutrophil transit through narrow spaces. This contrasts with the sADF setup, where large-scale nuclear remodeling is required to align the nucleus with the substrate. Together, these findings suggest that transient, small-scale nuclear constrictions in the C^3^ device are governed by local NE curvature changes rather than global nuclear shape remodeling.

After emerging from 3-micron constrictions, *cPLA_2_α* KO neutrophils exhibit significantly reduced migration distance, speed, and directionality compared to Scr and GFP-cPLA_2_α expressing neutrophils, underscoring a critical role for cPLA_2_α in post-constriction migratory performance. Notably, re-expression of GFP-cPLA_2_α in *cPLA_2_α* KO neutrophils restored this behavior, but the rescue was reversed when LTB_4_ synthesis was pharmacologically inhibited, highlighting that the functional contribution of cPLA_2_α depends on its catalytic role in LTB_4_ production. We previously demonstrated that LTB_4_ signaling mediates MLC II phosphorylation and drives cell polarization and directional migration toward fMLF (*22*). Here, we show that phosphorylation of MLC II at the cell cortex following 3 μm nuclear constriction is absent in *cPLA₂α* KO neutrophils, leading to impaired cell polarity and compromised chemotactic persistence. These findings support a model in which nuclear squeezing facilitates cPLA₂α enrichment at curvature-sensitive regions of the INM, facilitating its incorporation into NE-MVBs and subsequently into LTB_4_-producing exosomes. The released LTB_4_ then binds to LTB_4_ GPCRs and acts in an autocrine or paracrine manner to promote MLC II phosphorylation and sustain directional migration. Given the pivotal role of LTB_4_ in amplifying chemotactic signaling, this mechanism offers insight into the rapid, directed migration of neutrophils *in vivo*, where cells must traverse dense, collagen-rich tissue to reach sites of saturating chemoattractant concentrations (*6*). Interestingly, we also observed a striking morphological transition, from a pseudopod-driven amoeboid mode to a fan-shaped, keratocyte-like migration pattern, exclusively when GFP-cPLA_2_ neutrophils passed through 3-micron constrictions. In *Dictyostelium discoideum*, both amoeboid and keratocyte-like migration modes coexist and are dynamically regulated by external cues (*37*, *54*). Cells in the keratocyte-like state typically adopt a broad, laterally expanded lamellipodium with strong frontal actin protrusions and stable, rear-localized actomyosin (Myo II) contractility (*36*, *55*, *56*). This migration mode is supported by moderate substrate adhesion, similar to what is observed during neutrophil infiltration *in vivo* (*6*, *57*), whereas lower or higher adhesion levels tend to favor amoeboid or slower mesenchymal migration modes, respectively (*5*, *58*).

Ceramide and its metabolite ceramide-1-phosphate (C1P) have been shown to enhance cPLA₂α membrane recruitment and catalytic activity by interacting with its calcium-dependent C2 domain (*59*, *60*). This provides a mechanistic basis for the enrichment of cPLA₂α at ceramide-rich, lipid-ordered microdomains within the INM, where nSMase1 localizes and NE budding is initiated (*15*). Although cPLA₂α is present in NE-derived exosomes, its absence does not impact the number of 5LO/FLAP-positive exosomes. However, neutrophils expressing GFP-cPLA₂α – at higher levels than endogenous cPLA₂α in Scr cells – showed increased exosomal cPLA₂α and LTB_4_ levels, and exhibited enhanced post-constriction directional migration. These observations suggest that AA release by cPLA₂α is the rate-limiting step in exosomal LTB_4_ biosynthesis. Trypsin sensitivity of exosome-associated cPLA₂α indicates that its exposure on the exofacial surface aligns with its INM localization in migrating neutrophils. We propose that upon chemoattractant stimulation, the nuclear pool of cPLA₂α, capable of sensing high positive curvature, is selectively recruited to nascent intraluminal vesicles budding from the INM with NE-MVBs (**Fig. 7E**). These cPLA_2_α-containing exosomes then serve as a local hub for rapid LTB_4_ production and secretion. This model is supported by previous findings linking nuclear cPLA₂α enrichment with AA release and LTB_4_ production in okadaic acid-treated macrophages (*61*) and phosphatase-treated cancer cells (*62*), highlighting a conserved role for nuclear cPLA_2_α in curvature sensing and eicosanoid signaling. While the precise functions of cytoplasmic cPLA₂α remain to be fully defined, it has been implicated in prostaglandin synthesis – a distinct eicosanoid class – on lipid droplets derived from the tubular endoplasmic reticulum (*63*).

Together, our data suggest a novel function for nuclear cPLA₂α in establishing neutrophil polarity and promoting a transition towards keratocyte-like migration. This switch appears to be tightly linked to localized LTB_4_ production and downstream phosphorylation of MLC II, a key effector of MyoII activity. Given that rear MyoII contractility is essential for maintaining front-rear polarity and directional persistence, we propose that the cPLA₂α–LTB_4_–MLC II signaling axis acts as a mechano-chemical feedback loop that translates transient nuclear deformation into stable cell polarity and efficient post-constriction chemotaxis. These findings highlight a previously unrecognized mechanism by which confined migration triggers an exosome-mediated signaling cascade to reinforce migratory behavior. Future studies will help dissect how this axis governs migratory mode transitions in immune cells across different tissue contexts.

## ACKNOWLEDGMENTS

We are thankful to the past and present members of the Parent laboratory for advice and support. We acknowledge Dr. Michael Holistat and Amanda Prieur from the Platelet Pharmacology and Physiology Core at the University of Michigan for providing human blood from healthy volunteers and Peilin Shen for neutrophil isolation and technical assistance.

## Funding

National Institute of Health grant # EB030474 (BMB)

National Institute of Health grant #R01AI152517 (CAP)

National Institute of Health T32 Training Program in Cell and Molecular biology GM145470 (SPC)

American Heart Association pre-doctoral fellowship #014194 (FJJ)

American Heart Association pre-doctoral fellowship #025905 (SPC)

American Heart Association post-doctoral fellowship #916874 (SBA)

The funders had no role in study design, data collection and analysis, decision to publish, or preparation of the manuscript.

## Author contributions

Conceptualization: CAP, SBA, FJ-J. Designed and performed experiments: SBA and FJ-J. Generated new reagents/analytic tools: KL and BMB. Generated nanofibers: KL and BMB. Fabricated C^3^: YC and EY. Wrote Matlab code for chemotaxis analysis: SPC and LEH. Analyzed data: SBA, FJ-J. and SPC. Writing – original draft: SBA and CAP. Writing – review and editing: SBA, FJ-J, and CAP.

## Data and materials availability

All data needed to evaluate the conclusions in the paper are present in the paper and/or the Supplementary Materials. Raw microscopy images and source data for images are available from the corresponding author upon reasonable request. All CellProfiler pipelines used in this study are made available on the CellProfiler website linked to the publication weblink and accession information, to ensure transparency and reproducibility of the analysis. The Matlab code used to analyze underagarose migration of neutrophils has been uploaded to a publicly available repository and can be accessed at: Collie, S., Hein, L., Loesel, K., & Parent, C. underagarose-analysis (Version 1.0.2). Github. 2025. https://doi.org/10.5281/zenodo.15080547. Requests for reagents and resources should be directed to the lead contact, Carole Parent.

## SUPPLEMENTARY MATERIALS

### Methods and materials

#### Cell culture

The human myeloid leukemia-derived pro-myelocytic cell line HL60 was obtained from ATCC (CCL-240) and maintained in RPMI-1640 (Gibco 11875-093) medium containing 10% heat-inactivated FBS, 20 mM HEPES pH 7.2, and 100 U/ml penicillin-streptomycin antibiotic cocktail (ThermoFisher Scientific #15-140-122). To generate neutrophil-like cells, HL60 cells were differentiated in culture medium containing 1.3% DMSO for 6 days with a change to fresh medium every other day as described by Saunders *et al.* (*64*)

HEK293T cells obtained from ATCC (CRL-3216), cultured in DMEM supplemented with 10% FBS, were used to generate lentiviral particles for gene KO expression in HL60 cell lines. Lentiviral packaging plasmids pVSVG and psPax2, along with pLentiCRISPR V2 vector expressing either Scr single guide RNA (sgRNA), cPLA_2_α sgRNA, or pCDH MSCV MCS EF1 neomycin vector expressing cPLA_2_α fused with eGFP in-frame at its N-terminal, were transfected to HEK293T at a ratio of 1:2:4 using Lipofectamine 3000 transfection reagent. The culture supernatant containing lentiviral particles was collected after 48 h and 72 h post-transfection and pooled. Lentiviral particles were concentrated by incubating supernatants in 1x lentivirus concentrator (4x stock, 40% w/v PEG-8000 and 1.2 M NaCl in 1x PBS) overnight at 4 °C and precipitated at 1500g for 45 min at 4 °C. The concentrated virus was resuspended in RPMI-1640 containing 8 μg/mL hexadimethrine bromide (polybrene) (Sigma Aldrich, H9268-5G) and added to HL60 cells. The clones expressing the construct were selected in 2 µg/mL puromycin (pLentiCRISPR V2) or 1 mg/mL G418 (pCDH MSCV MCS EF1), dilution plated for single cell cloning, and verified using western blotting and genetic sequencing.

#### Plasmid constructs

The cPLA_2_α sgRNA, ACACCACTACCGTAAACTTG, was cloned into the pLentiCRISPR V2 plasmid, which was a kind gift from the Zhang lab. pCDH-puro-GFP-cPLA_2_α construct was cloned using the NEB Gibson assembly kit (NEB E5510). cPLA_2_α was amplified from GenScript plasmid pCDNA3.1-cPLA2 (Clone ID Ohu19957) using 5’-ctgtacaagATGTCATTTATAGATCCTTACCAG-3’ and 5’-ccctcagcggccgcggatccTGCTTTGGGTTTACTTAGAAAC-3’ and GFP was amplified from FPR1-eGFP plasmid from Subramanian *et al*. (*22*) using 5’-gagctagagctagcgaattcGCCACCATGGTGAGCAAG-3’ and 5’-taaatgacatCTTGTACAGCTCGTCCATGC-3’ primers. GFP-cPLA_2_α was amplified from pCDH-puro-GFP-cPLA_2_α construct using 5’-gcgggcGCTAGCATGGTGAGCAAGGGCGAGG-3’ and 5’-gcgcggcGCGGCCGCctaTGCTTTGGGTTTACTTAG-3’ primers and cloned into the NheI and NotI sites in pCDH MSCV MCS EF1 neomycin vector. All cloned constructs were confirmed by Sanger sequencing.

#### Isolation of human neutrophils

Blood was donated by healthy males and females who had not taken aspirin for seven days and NSAIDs for 48 hours. Blood was collected by venipuncture from the Platelet Pharmacology and Physiology Core at the University of Michigan. Neutrophils were purified using dextran-based sedimentation followed by histopaque density gradient centrifugation as described earlier (*27*). Briefly, whole blood was incubated with an equal volume of 3% dextran (Sigma D1037) in 0.9% NaCl for 1 h at 37 °C to sediment erythrocytes. One volume of Histopaque-1077 (Sigma 10771) was underlaid to three volumes of plasma containing monocytes, lymphocytes, and neutrophils and centrifuged at 400g for 20 min without brake to separate neutrophils from peripheral blood mononuclear cells (PBMCs). Residual RBCs were lysed using ACK lysis buffer (Gibco A10492-01, 100mL). Isolated neutrophils are resuspended in mHBSS (150mM NaCl, 4mM KCl, 1.2mM MgCl_2_, 5 mM glucose, and 20mM HEPES pH 7.2). This protocol yields >95% neutrophils.

#### Under Agarose chemotaxis assay and chemotaxis analysis

Chemotaxis assay was performed as previously described (*64*). Briefly, 0.5% SeaKem ME agarose (Lonza 50010) in 1:1 DPBS (Gibco 14190-144) and 1x mHBSS was solidified in either 35-mm glass bottom (Celvis) or 8-well chamber glass bottom chambers (Celvis C8-1.5H-N) pre-coated with 1% BSA (Sigma A7979-50mL) in DPBS. Two wells of 1 mm diameter each were carved 2 mm apart. Differentiated HL-60 cells were resuspended in washed in DPBS, counted, resuspended at a density of 10 million per ml, and stained using 1 μg/ml Hoechst 33342 nuclear stain (Invitrogen H21492) for 15 min at 37 °C at 10 RPM. 100 nM fMLF diluted in 1x mHBSS was added to one well, and 50,000 stained cells in 5 μL mHBSS were added to another well. Time-lapse images were acquired at 30-second intervals for 1.5 hours using a 10X objective of a fluorescent microscope equipped with an environment-controlled unit set at 37 °C.

Using the TrackMate plugin, all the migrating cells (200–1000) were tracked on the Hoechst 33342 channel. Spot statistics were downloaded from TrackMate analysis and uploaded into the chemotaxis analysis code in MATLAB R2021a. Tracks with less than 100 μm final distance on the x-axis were excluded from the final analysis. The average of chemotaxis parameters for each experiment was plotted using GraphPad Prism.

#### Expansion Microscopy and Immunofluorescence Staining

dHL-60 cells or PMNs were allowed to migrate under agarose towards 100 nM fMLF for 1 h and fixed with 4% paraformaldehyde (PFA) (Electron Microscopy Sciences, 15174) diluted in 1x mHBSS at 37 °C for 20 min. The agarose was scooped out, and the cells were fixed and blocked in staining solution (0.5% saponin and 2% goat serum in PBS) for 1 h at room temperature. Cells were stained overnight at 4 °C using primary antibodies diluted in a staining solution. The antibodies and their dilutions used for normal immunofluorescence staining are rabbit anti-GFP (1:2000, ThermoFisher A6455), rabbit anti-LBR (1:500, Abcam 32535), rabbit anti-FLAP (1 μg/ml, Abcam 85227), mouse anti-cPLA_2_α (1:100, Santacruz biotechnology 376618), rabbit anti-5 lipoxygenase (1:200: Abcam 169755), and rabbit anti-phospho MLC II (1:100, Cell Signaling Technology 3674S). Stained cells were washed once with 0.2% saponin in DPBS, followed by DPBS twice for 5 mins each, followed by incubation with AlexFluor-tagged secondary antibodies (1:500) in staining solution for 1 hour at room temperature. The cells were washed with DPBS thrice, overlaid with 150 μl ImmuMount™ mounting media (ThermoScientific, 9990402), and stored at 4 °C until imaging on a Zeiss 880 confocal/airyscan microscope.

For 4x expansion microscopy, GFP–cPLA2α dHL60 cells were allowed to migrate for 1 h at 37 °C under agarose on a BSA-coated 22 × 22 mm coverslip (#1.5) placed in a 35-mm dish. Cells were fixed with 1 mL of 4% paraformaldehyde and 0.05% glutaraldehyde in PHEM buffer (60 mM PIPES, 25 mM HEPES, 10 mM EGTA, 2 mM MgCl₂, pH 6.9) at 37 °C for 20 min. After gently removing the agarose, the coverslip-containing dish was incubated with 2 mL of anchoring solution (2% acrylamide and 1.4% formaldehyde in PBS) for 3 h at 37 °C. For gelation, the coverslip was removed from the gelation chamber, placed in a 35-mm dish, and incubated with 2 mL of denaturation buffer (200 mM SDS, 200 mM NaCl, 50 mM Tris base, pH 6.8) with shaking at room temperature until the gel detached, followed by denaturation at 85 °C for 90 min. The gel was expanded in double-distilled water (10 mL, five times) by incubating it at room temperature with shaking in a 10-cm dish for 20 min each time. The gel was then incubated in PBS containing Hoechst 33342 to visualize cells. Gel pieces (∼1 cm²) containing migrated cells were excised and transferred to 1.5 mL tubes containing 250 μL of antibody staining solution (PBS with 2% BSA) and the following primary antibodies: mouse anti-calnexin (1:50, Proteintech 66903-1-Ig), mouse anti-cPLA2α (1:50, Santacruz biotechnology 376618), rabbit anti-GFP (1:100, abcam 290), and rabbit anti-LBR (1:50, Abcam 32535). Incubation was performed overnight at 4 °C with end-over-end rotation at 10 RPM. Gels were washed with 500 μL PBS containing 0.1% Tween-20 three times for 10 min each with rotation. Secondary antibody incubation was carried out for 6 h at room temperature, followed by three washes with PBS + 0.1% Tween-20 and a final PBS wash. The gel was re-expanded in double-distilled water three times at room temperature for 20 min each, followed by overnight expansion at 4 °C to reach saturation. The expanded gel was immobilized (cell side down) on a poly-L-lysine–coated 12-mm glass-bottom 35-mm dish. Gels were gently pressed with a soft brush to prevent drift, surrounded with 100 μL double-distilled water to prevent shrinkage, and overlaid with a 22 × 22 mm coverslip to stabilize the sample. Imaging was performed using a Zeiss LSM 880 with Airyscan and a 63× (1.4 NA) oil objective. The effective resolution post-expansion was ∼30–40 nm laterally and ∼100 nm on the z-axis.

#### Intact nuclei isolation and immunostaining

This protocol is modified from the DRM isolation protocols by Persaud-Sawin *et al.* and Cascianelli *et al.* (*65*, *66*). Briefly, 50 million dHL60 were resuspended in 1 mL 1x mHBSS. To inhibit proteases, cells were pre-treated with 2 mM AEBSF hydrochloride (Pefabloc, Fisher Scientific AC328110010) for 15 min at 37 °C, before stimulation with 100 nM fMLF for either 15 or 30 min at 37 °C while rotating at 10 RPM. Cells were pelleted at 6,000*g* for 30 seconds, and the proteins were crosslinked using 10 mM dimethyl pimelimidate (DMP, TCL chemicals D4476) in 1x mHBSS for 15 min at 37 °C, followed by quenching free DMP using 20 mM Tris-Cl pH 8.0 for 10 min at room temperature. The plasma membrane was partially lysed twice to separate nuclei and cytosol by triturating cells (50 million/ml) 10 times in ice-cold hypotonic lysis buffer (10 mM HEPES pH 7.2, 4 mM MgCl_2_, 25 mM NaCl, 1 mM DTT, and 0.1% NP-40), using 1 ml pipette at a density of 50 million cells/mL followed by centrifugation until speed reaches 16,000g. Supernatants from the first lysis were collected as the cytosolic fraction. The pellet after the second centrifugation was washed twice with ice-cold Barnes solution (85 mM KCl, 85 mM NaCl, 2.5 mM MgCl_2_, and 5 mM trichloroacetic acid, pH 7.2) to remove residual ER fragments (microsomes) from the nuclei. Supernatants from the first wash were collected as the Barnes (ER) fraction. For the western blot analysis, all fractions were resuspended in 1X XT sample buffer, boiled at 95 °C for 10 min, and loaded in equal volume. For immunofluorescent staining, purified nuclei corresponding to that of 1 million cells were resuspended in 1 ml of ice-cold nuclei resuspension buffer (10 mM HEPES pH 7.2, 4 mM MgCl_2_, 150 mM NaCl, 1 mM DTT, and 250 mM sucrose) and added to poly L-Lysine (Sigma, P4832) coated #1.5 glass coverslips and centrifuged at 500*g* for 5 min at 4 °C. The isolated nuclei were then fixed using 4% PFA in resuspension buffer for 10 min at room temperature, followed by primary antibody staining overnight at 4 °C in DPBS containing 2% goat serum. The antibodies used are mouse anti-Ceramide (1:200, Sigma C8104-50TST), rabbit anti-GFP (1:500, ThermoFisher A6455), and rabbit anti-FLAP (1:200, Novus NB300-891).

#### Nuclear DRM extraction

For lipid-ordered microdomains extraction, membranes of ∼50 million isolated nuclei were solubilized in 650 μl ice-cold TNE buffer (50 mM Tris-Cl pH 7.4, 150 mM NaCl, and 5 mM EGTA) containing 4 mM MgCl_2_ and 1% Triton X-100. The suspension was homogenized by passing through a 23G needle 30 times, and the homogenate was incubated on ice for 30 min. Supernatants containing solubilized nuclear membranes and nucleoplasm were collected at 110g for 10 min at 4 °C, and the pellet (chromatin/nucleoskeleton) was discarded. Supernatants were adjusted to 40% optiprep using 60% OptiPrep stock solution (Sigma D1556) to a final volume of 2 ml and overlayed with 7mL of 30% iodixanol, 2mL of 20% iodixanol followed by 1mL of 5% iodixanol solution in TNE buffer, in 13 ml ultracentrifuge tubes (Beckman 331372). After centrifugation at 150,000g for 16 h at 4 °C, sixteen 750 µl fractions from the top were collected. The proteins in collected fraction were precipitated using trichloroacetic acid (TCA)-acetone method, air dried and were resuspended in 80 µl of 1x XT sample buffer (BioRad, 1610791) under reducing conditions, boiled at 95 °C for 10 min and loaded on Criterion XT 4-12% Bis-Tris gel (BioRad 3450124) for electrophoresis. The electrophoresed proteins were transferred to 0.2 μm PVDF membrane, blocked using 1X Fish gelatin (Fisher Scientific NC0382999) in Tris-buffered saline containing 0.1% Tween-20 (Fisher Scientific 337-500) and probed for specific proteins using antibody against FLAP (1 µg/mL, Abcam 85227), flotillin 2 (1:1,000, CST 3436), nSMase1 (1:500, CST 3867) and cPLA_2_α (1:1,000, Santa Cruz Biotechnology sc-376618), using electrochemiluminescence (ECL) capture on photographic films.

#### Generation of sADF fibslips

Dextran vinyl sulfone (DexVS) fiber mats were synthesized using electrospin technology as described previously by Loesel *et al*. (*32*). Briefly, 0.6 mg/ml dextran vinyl sulfone (DexVS) dissolved in dimethylformamide (DMF): MilliQ water (1:1) was mixed with 100 mg/mL lithium phenyl-2,4,6-trimethylbenzoylphosphinate (LAP), 0.75 mM methacryloxyethyl thiocarbamoyl rhodamine B, and 5 vol% glycidyl methacrylate to generate an electrospinning solution. DexVS solution was electrospun in a humidity-controlled glove box at 30% to 35% relative humidity and 0.2 ml/h flow rate. To create aligned fibers, an 18 mm^2^ glass coverslip (Fisher Scientific, 12546) was placed between two parallel copper electrodes set to –4.0 kV. The stainless-steel needle containing the polymer solution was situated 7 cm from the collection surface and connected to the voltage source set to +4.0 kV. Electrospun fibers were deposited onto the coverslip for 5 min to achieve the desired thickness of the fiber mat. The fiber mats were primarily crosslinked under ultraviolet light at 100 mW/cm^2^ for 20 seconds to stabilize the fibers. The fiber-coated coverslips were glued to the bottom of a modified 12-well plate using SYLGARD^TM^ 164 Silicone Elastomer kit (Dow, 0.4028273). The 12-well plates (Fisher Scientific, FB012928) were modified by drilling 15 mm^2^ holes at the bottom of a polystyrene 12-well plate using a Dremel 7760 tool.

Before the experiment, the fiber mats were functionalized with 2.5% (w/v) heparin methacrylate dissolved in 1 mg/mL LAP solution using 100mW/cm2 ultraviolet light exposure for 20 s. The 12-well plates are sterilized using 70% ethanol for 10 min, followed by coating with 10 μg/mL fibrinogen (Sigma, 4129) in DPBS for 1 h at 37 °C, before plating cells.

#### Constricted chemotaxis chamber (C^3^) fabrication

We first generated a computational model of C^3^ using AutoCAD (Autodesk). The master molds were then fabricated employing the standard photolithography method using SU-8 photoresist (MicroChem) following the manufacturer’s protocol. Two masks were used to fabricate the multiple heights, one for the migration channels (5 μm high) and the other for the cell/chemoattractant channels (100 μm high). Poly-dimethylsiloxane (PDMS, Sylgard 184, Dow Corning) was prepared with the 10:1 elastomer-to-curing agent ratio. PDMS was poured on the channel molds and cured at 80 °C for 4 h to polymerize, before peeling to generate a PDMS layer of C^3^. The PDMS layer and the substrate (#1.5 glass coverslip 22×22 mm) were activated by oxygen plasma treatment to facilitate bonding. The devices are stored at room temperature in a moisture-free environment. Before the experiment, the C^3^ was desiccated at 150 °C for 5 min and sterilized for 20 min by UV exposure in a laminar flow hood. C^3^ channels and migration chamber were then filled with 1% BSA in DPBS and incubated for 1 h at 37 °C to coat the glass and PDMS surface and avoid excessive cell adhesion on negatively charged surfaces, which impedes migration. The channels were then washed with DPBS and then filled with 1x mHBSS for the experiment. Ten μl of dHL60 cell (Hoechst 33342-stained) suspension at a density of 20 million cells per ml was added to the cell inlet port, and 10 μl of chemoattractant (100 nM fMLF in mHBSS) was added to chemoattractant inlet port. Three microliters of cell suspension and chemoattractant each were aspirated from their respective outlets. The cells were allowed to chemotax towards increasing chemoattractant concentration in the migration chamber and imaged using a 10x objective on a Zeiss Colibri microscope fitted with an environment control chamber maintained at 37 °C and >90% humidity. Image acquisition was automated using MetaMorph software.

#### Microscopy and Image Analysis

Fixed and immunostained cells/isolated nuclei were imaged using the Plan Apochromat 63X/1.4 Oil DIC M27 objective on Zeiss LSM 880, AxioObserver equipped with AiryScan Superresolution mode. For colocalization analysis, ROIs of isolated nuclei were thresholded using maximum entropy parameters, and Mander’s co-localization coefficients were determined using the Coloc2 analysis plugin in FIJI. The values were plotted using GraphPad Prism software. Outliers were calculated using the GraphPad Prism ROUT test (Q=1%) and were excluded from the final statistical analysis using the Mann-Whitney test.

#### Exosome isolation, LTB_4_ ELISA, and trypsin protection assay

Exosome isolation was performed according to the guidelines described by Welsh *et al*. (*67*). Scr, *cPLA_2_α* KO, and GFP-cPLA_2_α dHL60 neutrophils were stimulated with 100 nM fMLF in 1x mHBSS containing 10 Units/ml DNase I (Sigma Aldrich DN25) for 30 min at 37 °C. The cells were pelleted at 500*g* for 5 min at 4 °C, and supernatants were collected and centrifuged again at 4000*g* for 20 min to remove microvesicles and apoptotic bodies. The supernatant was mixed 1:1 with 16% PEG-6000 (Bio Basic PB0432) dissolved in 20 mM HEPES pH 6.9 and 500 mM NaCl to precipitate extracellular vesicles at 4 °C for 36 h, followed by centrifugation at 4,000*g* at 4 °C for 1 h. The PEG-precipitated EVs were washed by resuspending the pellet in 5 mL ice-cold PBS and centrifugation at 100,000*g* for 1 h at 4 °C in the Beckman SW55Ti rotor. The concentrated extracellular vesicles were resuspended in 1 ml of 250 mM sucrose and 20 mM Tris-Cl pH 7.4, overlayed on top of optiprep gradients, and centrifuged at 100,000*g* for 16 h at 4°C using a Beckman Sw41Ti rotor. The optiprep gradients prepared in 250 mM sucrose and 20 mM Tris-Cl pH 7.4, were layered starting from the bottom as 3 ml of 40% optiprep, 3 ml of 20% optiprep, 3 ml of 10% optiprep, and 2 ml of 5% optiprep. The fractionated exosomes were collected as 12 fractions of 1 ml each, starting from the top (lower to higher density). Exosome-enriched fractions 4-9 (Iodixanol density 1.083-1.142 g/ml) were pooled and diluted to 13 ml with PBS, followed by centrifugation at 100,000*g* for 1 h at 4 °C. The purified exosomes were used to assess exosomal LTB_4_ content and determine the distribution of exosome-associated proteins.

LTB_4_ ELISA kit (Cayman Chemicals 520111) was used to assess LTB_4_ levels within the isolated exosomes homogenized in 100 μl ELISA buffer using a 3 mm diameter sonicator probe at an amplitude of 20% with 2 sec on/off cycles for a total of ten cycles on ice. To detect LTB_4_ concentrations within the linear range, 50 μl of concentrated homogenate was diluted 4x in ELISA buffer, and LTB_4_ levels were quantified according to the manufacturer’s instructions. The values obtained were plotted using GraphPad Prism.

To determine the distribution of proteins within the exosomes, the isolated exosomes were resuspended in mHBSS supplemented with 1mM CaCl_2_, volumetrically divided into two equal fractions, and one fraction treated with 50 µg/ml trypsin (ThermoFisher Scientific 25200072) for 30 min at 37 °C. The trypsin was inactivated by diluting the exosomes in mHBSS supplemented with AEBSF hydrochloride, followed by centrifugation at 120,000*g* for 1 h at 4 °C to pellet the exosomes. The pelleted exosomes were lysed in 1x XT sample buffer at 95 °C for 10 min, and equal volumes were loaded on a Criterion XT 4-12% Bis-Tris electrophoresis gel for western blotting.

#### Generation of whole cell lysate for western blotting

dHL60 neutrophils were washed once with DPBS, resuspended in mHBSS, and treated with 2 mM Pefabloc™ for 15 min while rotating. Cells were pelleted at 6000*g* for 30 sec and resuspended in 1x Laemmli sample buffer (Fisher Scientific AAJ61337AD) diluted in PBS. Samples were boiled at 95 °C for 10 min, and lysate volume equivalent to 250,000 cells was loaded on a 4-20% Tris-Glycine gel (Invitrogen XP04205BOX) for electrophoresis. The proteins were transferred to 0.2 μm nitrocellulose membrane (MDI, SCNX8401XXXX101), blocked using 1x Fish gelatin (Fisher Scientific NC0382999) in 0.1% Tween-20 containing Tris buffered saline, and probed for specific proteins using antibody against cPLA_2_α (1:1,000, Santa Cruz Biotechnology sc-376618), FLAP (1 µg ml^−1^, Abcam 85227), 5LO (1:1000, Abcam, ab169755), LTA_4_H (1:1000 Protein Tech 13662-1-AP) and GAPDH (1:1000, Santa Cruz Biotechnology).

#### Statistics and reproducibility

All data presented here are from at least three independent biological replicates. Appropriate significance tests have been used to determine the level of confidence and variability in the data mentioned in the figure legends.

**Fig. S1.**
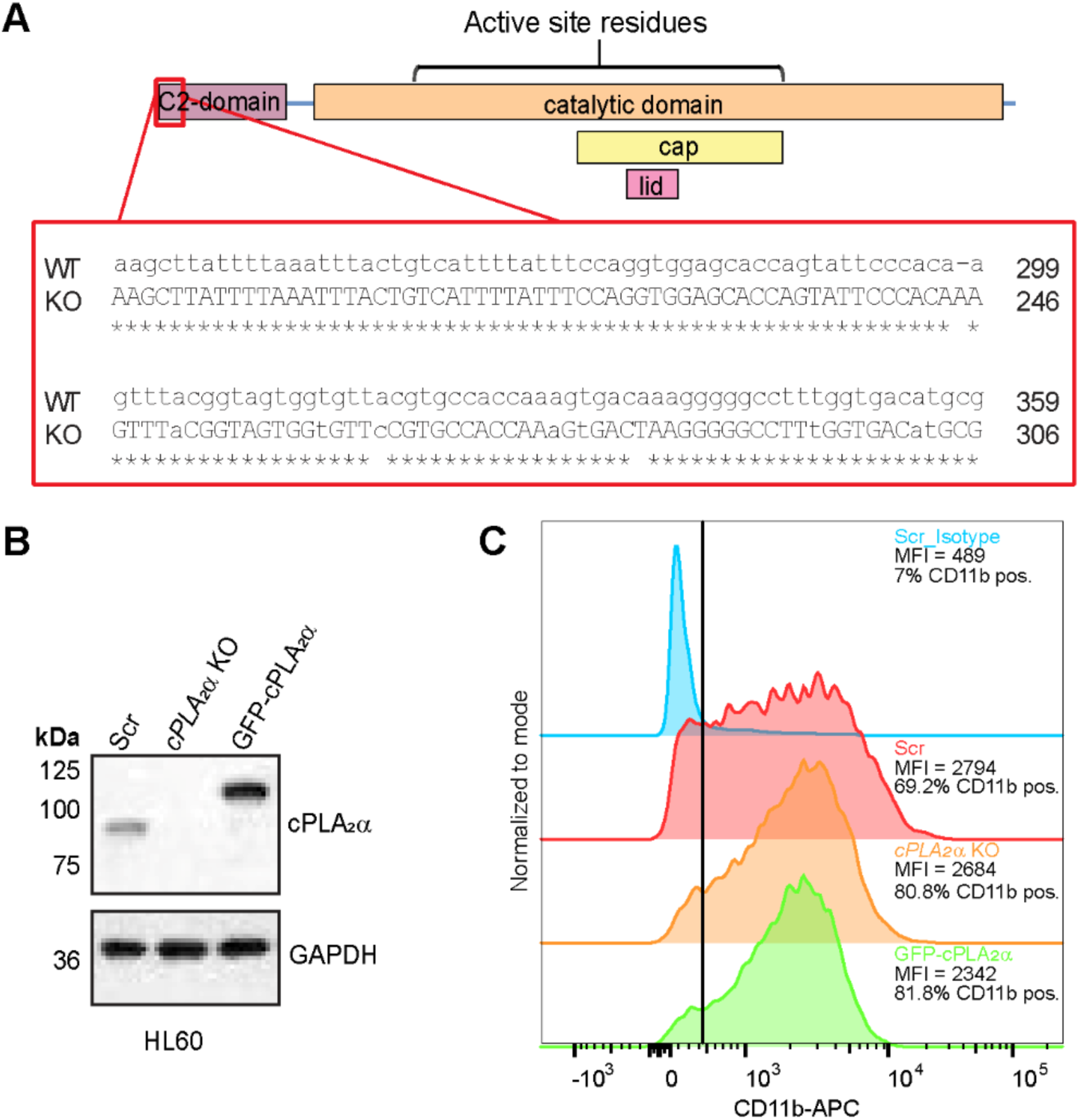
Validation of cPLA_2_α genetic manipulation. **(A)** Schematic showing the mutations introduced at the beginning of the cPLA_2_α C2-domain using the CRISPR-cas9 approach. (**B)** Immunoblot of HL60 cells showing the level of cPLA_2_α expression in KO and rescue cells. (**C**) Histogram showing the percent CD11b-positive Scr, *cPLA_2_α* KO, and GFP-cPLA_2_α dHL60 cells, and levels of surface CD11b.

**Fig. S2.**
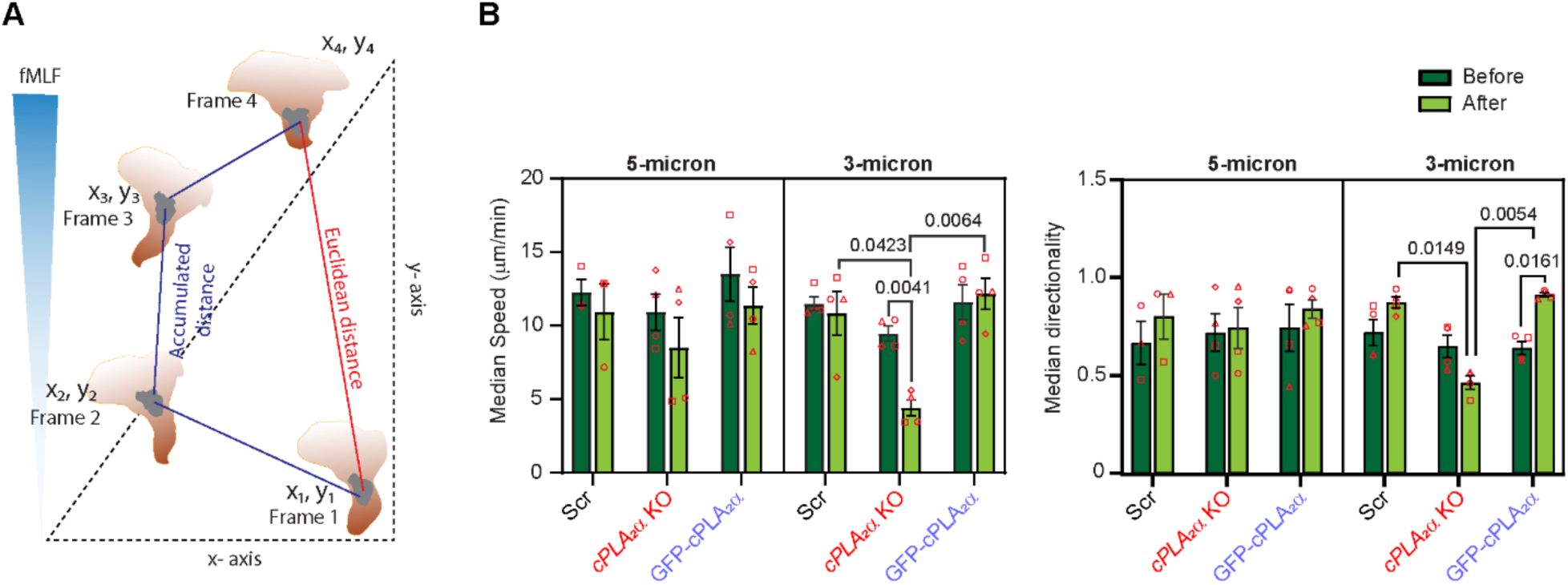
Measurement of chemotaxis parameters in C^3^. **(A)** Schematic showing the approach used to quantify chemotaxis parameters. **(B)** Graphs showing the changes in post-constriction median speed (left) and median directionality (right) of chemotaxing Scr, *cPLA_2_α* KO, and GFP-cPLA_2_α dHL60 cells, compared to pre-constriction parameters. N=4. Graphs are plotted as mean ± s.e.m. *P* values determined using three-way ANOVA are shown.

**Fig. S3.**
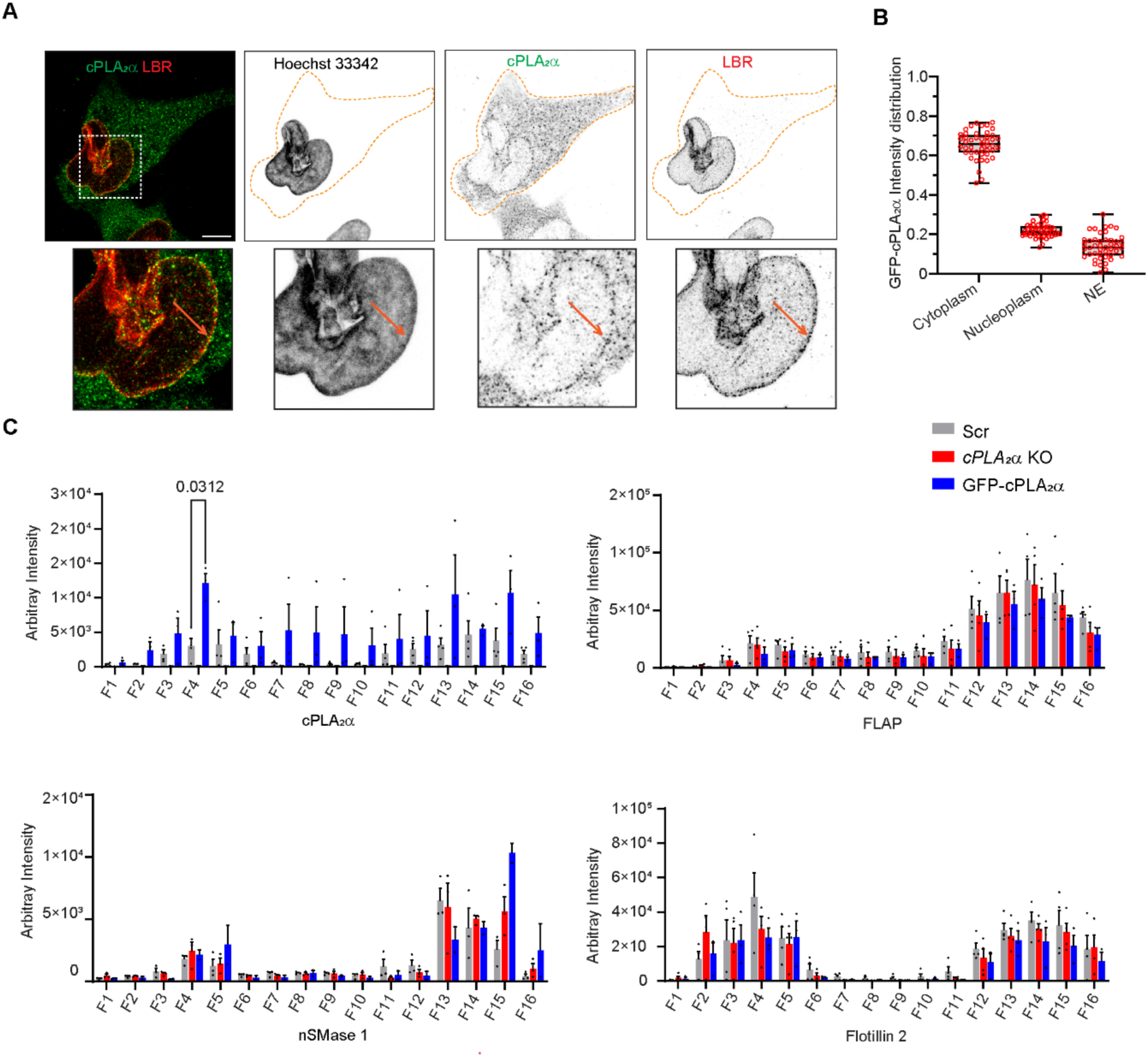
NE distribution of GFP-cPLA_2_α in dHL60 neutrophils. **(A-B)** Four-fold expansion microscopy images (A) of dHL60 neutrophils stably expressing GFP-cPLA_2_α chemotaxing towards fMLF, fixed and stained with anti-GFP antibody and LBR, and quantified (B) for cytoplasm, nucleoplasm, and NE distribution. Scale is 5 µm. Datapoints (45, red circles) from 3 independent experiments are plotted as mean ± s.e.m., and *P* values determined using ordinary one-way ANOVA are shown. **(C)** Graphs showing the cPLA_2_α, FLAP, nSMase 1, and Flotillin 2 distribution in DRM and DSM fractions of NE obtained from activated dHL60 neutrophils. Data are plotted as mean ± s.e.m. of 4 independent experiments, and *P* values determined using ratio-paired t-test are shown.

**Movie S1:** 3D volumetric view of the dHL60 neutrophils migrating under agarose towards fMLF, fixed and stained with phalloidin (red) and LBR (green). Representative of N=3.

**Movie S2:** 3D volumetric view of the fMLF-activated dHL60 neutrophils plated over aligned microfibers (red), fixed and stained for phalloidin (green) and Hoechst 33342 (blue). Representative of N=3.

**Movie S3:** Chemotaxis of GFP-cPLA_2_α expressing and *cPLA_2_α* KO dHL60 neutrophils, stained with Hoechst 33342 (magenta), migrating through 3-micron constriction C^3^. Frame rate is 3 fps. Representative of N=4.

**Movie S4:** 3D volumetric view of the isolated nuclei showing the increased association of GFP-cPLA_2_α with ceramide at the NE upon neutrophil activation. Representative of N=3.

**Methods only references: 63 to 67**

